# Basolateral Amygdala Projections to the Prelimbic Cortex Suppress the Motivational Value of Reward-Associated Cues

**DOI:** 10.1101/2025.11.06.687050

**Authors:** Mandy Rita LeCocq, Ana Tue, Domiziana Casale, Isabel Laplante, Karim Elrakhawy, Anne-Noël Samaha

## Abstract

Reward-associated cues drive reward-seeking behaviours, with different cue types taking on distinct associative meanings. In instrumental settings, discriminative stimuli (DSs) signal when a response will produce reward, while conditioned stimuli (CSs) are paired with reward delivery after responding. Basolateral amygdala (BLA) projections are known to mediate CS-guided reward seeking, but their role in DS-guided behaviour remains unclear despite DSs being uniquely effective in triggering reward seeking. Here, we used optogenetic stimulation of BLA projections to the nucleus accumbens (NAc) core or prelimbic cortex (PrL) to probe their causal involvement in DS- and CS-guided sucrose seeking. Female and male rats learned to lever press during DS+ trials to obtain sucrose paired with a CS+, and to withhold responding during DS-trials where sucrose was unavailable. We then assessed sucrose seeking evoked by response-independent DS+ and CS+ presentations, without sucrose. The DS+ prompted the greatest increases in lever pressing, acting as a strong Pavlovian conditioned excitor. Cue-paired photostimulation of BLA→NAc core or BLA→PrL projections did not alter this effect. In a test of instrumental responding for conditioned reinforcement, rats lever pressed most for DS+ and CS+ presentations, with no effect of cue-type, indicating that the two cues gained comparable conditioned reinforcing value. While cue-paired photostimulation of BLA→NAc core neurons had no impact, photostimulation of BLA→PrL projections suppressed instrumental responding for both the DS+ and CS+. These findings reveal the BLA→PrL pathway as a key regulator that constrains control over behaviour by reward-associated cues, making them less desirable and reducing their motivational effects.

## Introduction

Environmental cues that predict rewards guide advantageous behaviours—such as seeking safety, water, and food—thereby supporting survival. However, these same cues can also drive maladaptive behaviours, contributing to psychopathologies like binge eating, addiction, and gambling disorders [1–3]. Delineating the behavioural and neurobiological processes by which reward cues guide responding is critical to understanding both adaptive and maladaptive reward-seeking behaviour that arise when these processes go awry.

Reward-predictive cues can acquire distinct associative meanings. In instrumental settings, discriminative stimuli (DSs) are encountered independently of an animal’s actions and signal when a specific action will be reinforced, while conditioned stimuli (CSs) occur as a consequence of seeking and are paired with reward delivery [4–6]. For instance, the sound of a stream acts as a DS, signalling water availability before any seeking begins, whereas the sensation of cool water in an animal’s mouth functions as a CS as it follows drinking behaviour and predicts imminent reward delivery.

DSs and CSs exert similar but also distinct behavioural effects. Both DSs and CSs can maintain reward seeking by reinforcing instrumental responses, thereby bridging delays to reward [7–13]. However, when presented unexpectedly and independently of behaviour, DSs reliably trigger robust increases in reward seeking [12–21], while CSs typically do not [7,12–14,19–21]. Thus, DSs are especially potent motivators of reward pursuit.

The basolateral amygdala (BLA) is critical for processing reward-associated cues. Inhibiting the BLA suppresses both CS-based conditioned reinforcement of reward-seeking behaviours [22–24] and DS-triggered increases in reward seeking [19,25–27]. Within the BLA, local glutamate neurotransmission is also required for both CS- and DS-guided reward seeking [14,28–31]. Together, this evidence suggests that BLA-dependent pathways mediate the effects of both DSs and CSs on reward seeking.

In this context, two major BLA output pathways—BLA projections to the nucleus accumbens (BLA→NAc) and to the prelimbic cortex (BLA→PrL)—are of interest. Both the NAc and PrL receive dense glutamatergic inputs from the BLA [32] and both pathways facilitate instrumental responding for drug-paired CSs, supporting their role in conditioned reinforcement [33–36]. While the contribution of these specific BLA pathways in DS-guided reward-seeking behaviour remains unknown, both the NAc and PrL mediate DS-driven behaviours [25,27,37–41].

Here, we evaluated the hypothesis that activation of BLA→NAc and/or BLA→PrL pathways is sufficient to enhance the motivational impact of reward-associated DSs and CSs. Using in vivo optogenetics, we paired cue presentation with circuit-specific stimulation and assessed effects on i) increases in sucrose seeking triggered by response-independent DS or CS presentation, and ii) the ability of response-dependent DS and CS presentation to reinforce instrumental responding, in the absence of reward.

## Methodology

### Subjects

All procedures followed the Canadian Council on Animal Care guidelines and were approved by the Université de Montréal. Sprague-Dawley rats were single-housed on arrival (Experiment 1: *n* = 16 females; *n* = 15 males; Experiment 2: *n* = 15 females; *n* = 16 males; females 9 weeks old, males 7 weeks old; Charles River, Raleigh, North Carolina – RMS, Barrier R04). After an initial 72 hours, rats were handled daily by the experimenters. Cages contained sani-chip bedding, a nylabone (Bio-Serv, K3580), and unrestricted access to food (Charles River Laboratories, Saint-Constant, Quebec, Canada). Access to water was restricted as specified below. Cages were held in a climate-controlled colony room (22 ± 1°C, 30 ± 10% humidity) on a reverse 12 h light/dark cycle (8:30 AM lights OFF). All experiments were conducted during the dark phase.

Rats used in Experiment 1 were first used in an experiment involving optogenetic stimulation of BLA cell bodies and therefore had bilateral optic fibers implanted into the BLA. Two weeks before being used in Experiment 1, all rats received laser delivery into the BLA, during which 10/19 ChR2-expressing rats exhibited photostimulation-induced seizures. Compared to the 9 rats that did not exhibit seizures, these rats showed similar behavioural responses during all training and testing phases in Experiment 1 (See Table S1), therefore, they were kept in the data set.

### Apparatus

Behavioural procedures were conducted in conditioning chambers with standard grid floors enclosed in sound-attenuating, ventilated melamine cubicles (Med Associates Inc., St-Albans, VT). The back wall of each chamber held a sound generator (top right; 2900 Hz, 75 dB) and two cue lights (one top center, one top left). On the front wall, two retractable levers were located on either side of a magazine, and a cue light was located above each lever. During sucrose self-administration and discrimination training, presses on the active lever (counterbalanced between left- and right-side levers) produced 0.1 ml of 10% *w*/*v* liquid sucrose (Sucrose, Bioshop, Burlington O.N.; dissolved in tap water) delivered via a liquid dispenser connected to a 20-ml syringe. Interruption of infrared motion detectors at the top and bottom of the magazine opening measured magazine entries. A clicker generator was located outside of the chambers. Two rows of infrared beam sensors, mounted 2.5 cm above the grid floor of each conditioning chamber, measured horizontal locomotor activity. Med PC IV software on a PC delivered stimuli and recorded responses.

### Intracranial Surgery

Using validated methods [8, 42], we administered 1 µl (Experiment 1: bilateral; Experiment 2: unilateral) of an optically active or control virus into the BLA. Experimental rats (Experiment 1: *n* = 19, Experiment 2: *n* = 19) received an infusion of AAV_2/7_-CamKIIa-hChR2_(H134R)_-eYFP (1.5E13 GC/ml; provided by Dr. Karl Deisseroth; Canadian Neurophotonics Platform Viral Vector Core Facility, Quebec) into the BLA (AP: -2.8 mm, ML: ±5.0 mm, relative to Bregma; DV: -8.4 mm relative to skull surface). Control rats (Experiment 1: *n* = 12, Experiment 2: *n* = 12) received an optically inactive virus (AAV_2/7_-CamKIIa-eYFP; 1.5E13 GC/ml; provided by Ed Boyden; Molecular Tools, Quebec).

An optic fiber implant (made in house; 300 µm core diameter, numerical aperture 0.39; Thorlabs, Saint-Laurent, Q.C.; glued with epoxy to a ferrule, Fiber Instrument Sales, Oriskany, N.Y.) was implanted into either the NAc core (Experiment 1; AP: +1.9 mm, ML: ±1.6 mm, both relative to Bregma; DV: −6.7 mm relative to skull surface) or the PrL (Experiment 2; AP: +3.20, ML: ±0.6, both relative to Bregma; DV: -3.7 relative to skull surface) of the ipsilateral hemisphere. Optogenetic manipulations began at least 6 weeks following surgery to allow sufficient viral expression.

### c-Fos immunohistochemistry

We have previously confirmed that optical stimulation under our conditions evokes action potentials in ChR2-eYFP-expressing BLA→NAc core neurons [42]. Here, we used immunohistochemistry to confirm that optical stimulation of ChR2-eYFP-expressing BLA terminals in the PrL increases c-Fos expression levels in the PrL, as a marker of cell activity. Rats expressing ChR2-eYFP (*n* = 11) or eYFP (*n* = 8) received a 33-min session during which 15 trials of 5-s optical stimulation were delivered. Stimulation was delivered on a variable interval 125-s schedule without cue presentations, which matched the parameters used in the subsequent cue-evoked seeking tests. Rats returned to their home cage for 80 min before being perfused with phosphate-buffered saline then 4% paraformaldehyde. Coronal sections (40 µm) containing the PrL were collected for c-Fos immunohistochemistry using established procedures [20]. Following immunohistochemistry, a section between +3.70 to +2.70 mm AP relative to Bregma, in which the optical fiber track was visible, was identified for each rat and imaged (LAS X Software; Leica) at x10 magnification using a fluorescent microscope (Leica, DM2000) equipped with a camera (Leica DFC425 C). c-Fos positive cells were automatically counted using a set threshold for detection with Trainable Weka Segmentation Fiji plugins (ImageJ; National Institute of Health). c-Fos positive cells were counted in a 432 x 432 µm surface area directly below the optic fiber and averaged across 1 to 3 sections per animal. For each rat, we also quantified c-Fos levels in the hemisphere contralateral to the fiber-containing hemisphere, at the same coordinates and region of the PrL. This provided control levels of c-Fos protein expression, in the absence of optogenetic stimulation.

### General Procedures

#### Sucrose Habituation

Rats were first habituated to drinking the sucrose solution by receiving 48-h access to a bottle containing 80 ml of 10% w/v sucrose in their home cage in addition to their regular water bottle.

#### Water Restriction

The day following sucrose habituation, water restriction began to promote liquid sucrose self-administration. Rats had 6 hours/day of water access for the first 2 days, then 4 hours/day for 2 days, then 2 hours/day until the end of the experiment. Rats received access to water at least 1 h after behavioural training.

#### Magazine Training

Two days after sucrose habituation, rats received 2 magazine training sessions (30 min/session) to become familiar with receiving sucrose in the magazine. During each session, 0.1 ml of sucrose was delivered on a VI 45-s schedule, totaling 30 deliveries per session. At the end of each training session, the magazines were checked to verify that the sucrose was retrieved. Throughout the training phases, all rats consumed the sucrose.

#### Autoshaping

Rats can show significant individual variability in their response to reward-predictive CSs and DSs. Rats with a sign-tracker Pavlovian conditioned approach phenotype are thought to respond more to CSs, while goal-tracker phenotypes are thought to respond more to DSs [15,43,44; c.f., 45,46. As such, to eventually account for individual variability in responding to the DS and CS in the present study, we first identified the rats’ Pavlovian conditioned approach phenotype during autoshaping sessions (30 min/session; Experiment 1: 6 sessions; Experiment 2: 9 sessions).

Sessions began with the lever-CS extending for 8 s, followed by the delivery of 0.1 ml of sucrose solution. Lever-sucrose pairings occurred on a VI 30-s schedule with 30 pairings per session. The lever-CS was counterbalanced across right- and left-side levers, and this lever was designated the active lever in subsequent training phases. Upon lever-CS presentation, preferentially approaching the site of reward delivery (i.e., the magazine) is a goal-tracking response while preferentially approaching the lever is a sign-tracking response (Fig S1A).

A response bias score was calculated to identify the rats’ Pavlovian conditioned approach phenotype across sessions (Figs S1B-C), using data averaged over the last 2 autoshaping sessions [(CS lever presses - CS port entries) / (CS lever presses + CS port entries)]. A score of +0.50 or higher indicates sign-tracking, a score of -0.50 or lower indicates goal-tracking, and a score of -0.49 to +0.49 indicates intermediate responding [47,48].

#### Sucrose Self-Administration

Following autoshaping, rats learned to self-administer sucrose in daily sessions (30 min/session). Each session began with both levers extending. Pressing the active lever produced 0.1 ml of sucrose, and pressing the inactive lever had no programmed consequence. Initially, sessions were conducted under a fixed ratio one (FR1) reinforcement schedule. Once rats earned 20 reinforcers and made twice as many active vs. inactive lever presses on two consecutive sessions, they moved to an FR3 schedule. Once rats earned 30 reinforcers and made twice as many active vs. inactive lever presses under FR3, they moved to discrimination training. In Experiment 1, rats reached each of FR1 and FR3 acquisition criteria in 2 sessions. In Experiment 2, rats reached FR1 criteria in 2 sessions and FR3 criteria in 3 sessions (Fig S2A).

#### Responding for Sucrose under Progressive Ratio

In Experiment 2, we measured baseline motivation to obtain sucrose to determine whether this significantly correlated with active lever pressing during subsequent cue-evoked seeking and conditioned reinforcement tests. After sucrose self-administration training, rats received one more self-administration session where sucrose was available under a progressive ratio schedule of reinforcement according to the following formula [(round[5e(reinforcer number × 0.2) – 5]) [49]]. The session ended either 1 h after the last self-administered sucrose delivery or after 5 h. We defined breakpoint as the last completed ratio requirement before either of these two outcomes (Figs S2B-D).

#### Discrimination Training

##### DS presentations in predictable order

Following sucrose self-administration training, rats in both experiments initially received 9 discrimination training sessions (63 min/session) to learn that lever pressing produced sucrose when a DS+ was present, but not when a DS- was present. Fig 1A shows stimulus modalities. The DS+ was the cue light above the active or inactive lever (counterbalanced), and the DS- was the cue light at the top center of the back wall. During discrimination training sessions, the rats also learned that a CS+ was paired with self-administered sucrose while a CS- was never paired with sucrose. For half of the rats, the CS+ was the cue light above the active or inactive lever (counterbalanced) paired with a tone, and the CS-- was the cue light at the top left corner of the back wall paired with a clicker. For the other half, the CS+ and CS-modalities were reversed.

**Fig 1.**
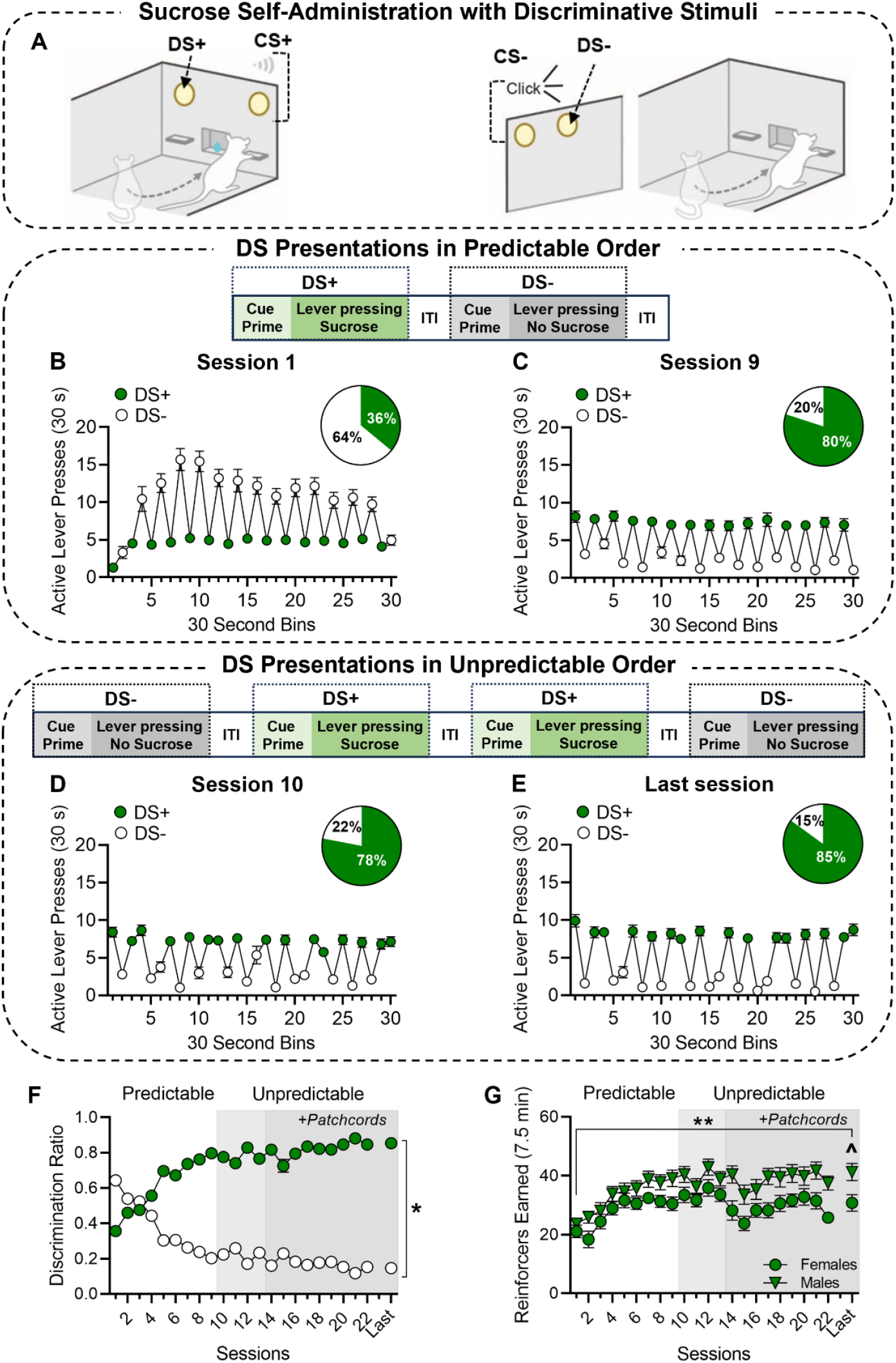
Sucrose self-administration comes under the control of discriminative stimuli. **(A)** Schematic of the discrimination procedure. Active lever presses made during DS+ (green circles) and DS- (white circles) trials on **(B)** Session 1 and **(C)** Session 9 where the DSs were presented in alternating, predictable order, and **(D)** Session 10 and **(E)** the last session where the DSs were presented in unpredictable order. Insets represent the percentage of active lever presses made during DS+ (green) vs. DS- (white) trials. **(F)** DS+ and DS-discrimination ratios and **(G)** sucrose reinforcers earned across sessions. In **(F)** and **(G)** light gray shading indicates sessions during which the DSs were presented in unpredictable order and darker gray shading indicates sessions where the rats’ optic fibers were also connected to patch cords. Data are average ± SEM. ‘Last Session’ represents combined data from the last session in each experiment (Session 20 in Experiment 1; Session 22 in Experiment 2). DS+, discriminative stimulus signaling sucrose availability. DS-, discriminative stimulus signaling sucrose unavailability. ITI, intertrial interval. * DS+ ratio > DS-ratio on Last session. ** Reinforcers earned on Session 1 > Last session in males. ^Males > Females on Last Session. *n* = 22 females, 23 males. **(A)** Adapted from Servonnet et al. (2023).

Sessions began with the levers retracted. After a 20 s pre-session interval, a DS+ cue prime was presented for 10 s. During the next 30 s, the DS+ remained on and both levers extended into the chamber. Active lever presses during these 30-s DS+ trials produced sucrose delivery under a random ratio 3 schedule. Each sucrose delivery was paired with a 5-s CS+ presentation. Lever pressing during CS+ presentations had no programmed consequence. The DS+ trial was followed by a VI 1.5 min intertrial interval (ITI), during which no cues were presented, and the levers were retracted. Next, a DS-cue prime was presented for 10 s, followed by a 30-s period during which the DS-remained on and the levers were extended into the chamber. During DS-trials, sucrose was not available and active lever presses produced a 5-s CS-presentation under a random ratio 3 schedule. During these initial 9 sessions, DS+ and DS-trials were presented in alternating, predictable sequence, until rats received 15 presentations of each DS type.

##### DS presentations in unpredictable order

To ensure that rats were assigning Pavlovian conditioned properties to the DSs, rather than keeping time to predict sucrose availability, we gave them additional discrimination sessions (Experiment 1: 11 sessions; Experiment 2: 13 sessions) during which the DSs were presented in unpredictable order. Sessions alternated daily such that they either began with a DS+ or DS-presentation, followed by pseudorandom presentations of the DS+ and DS-throughout the session. During the last 7 (Experiment 1) or 9 (Experiment 2) of these sessions, rats were tethered to patch cords (built in-house as described by [50]) without laser delivery to habituate them to subsequent optogenetic procedures.

A discrimination ratio was calculated to determine the ratio of active lever presses during DS+ vs. DS-presentations [DS+ ratio: (total DS+ active lever presses/total DS+ and DS-active lever presses); DS-ratio: (total DS-active lever presses/total DS+ and DS-active lever presses)]. Rats that had a DS+ ratio of ≥ 60% and earned ≥6 reinforcers/session, averaged across two consecutive sessions, were considered to have learned the discrimination task.

### Experiment 1. Effects of Optogenetic Stimulation of BLA →NAc core Neurons on Cue-Guided Sucrose Seeking

#### (DS+)-Evoked Increases in Sucrose Seeking

Here we tested the hypothesis that cue-paired BLA→NAc core neuron stimulation enhances the ability of response-independent presentation of the DS+ to trigger increases in sucrose-seeking behaviour. 24 h after the last discrimination training session, all rats received 2 cue-evoked seeking tests (33 min/session). As Fig 2A illustrates, sessions began with a 5-s priming period during which the DS+, CS+, DS-, CS-, or a ‘No Cue’ condition was presented response-independently, in counterbalanced order. This was followed by a 30-s lever access period during which the cue condition presented during the priming period remained on, and the levers were inserted. Pressing on either lever had no programmed consequence (no sucrose was delivered), and the rate of active lever pressing served as a measure of sucrose-seeking behaviour. A VI 1.5-min ITI followed, during which no stimuli were presented and both levers retracted. The ITI was followed by the next 5-s priming period and 30-s lever access period. This cycle repeated until each cue condition was presented 3 times.

**Fig 2.**
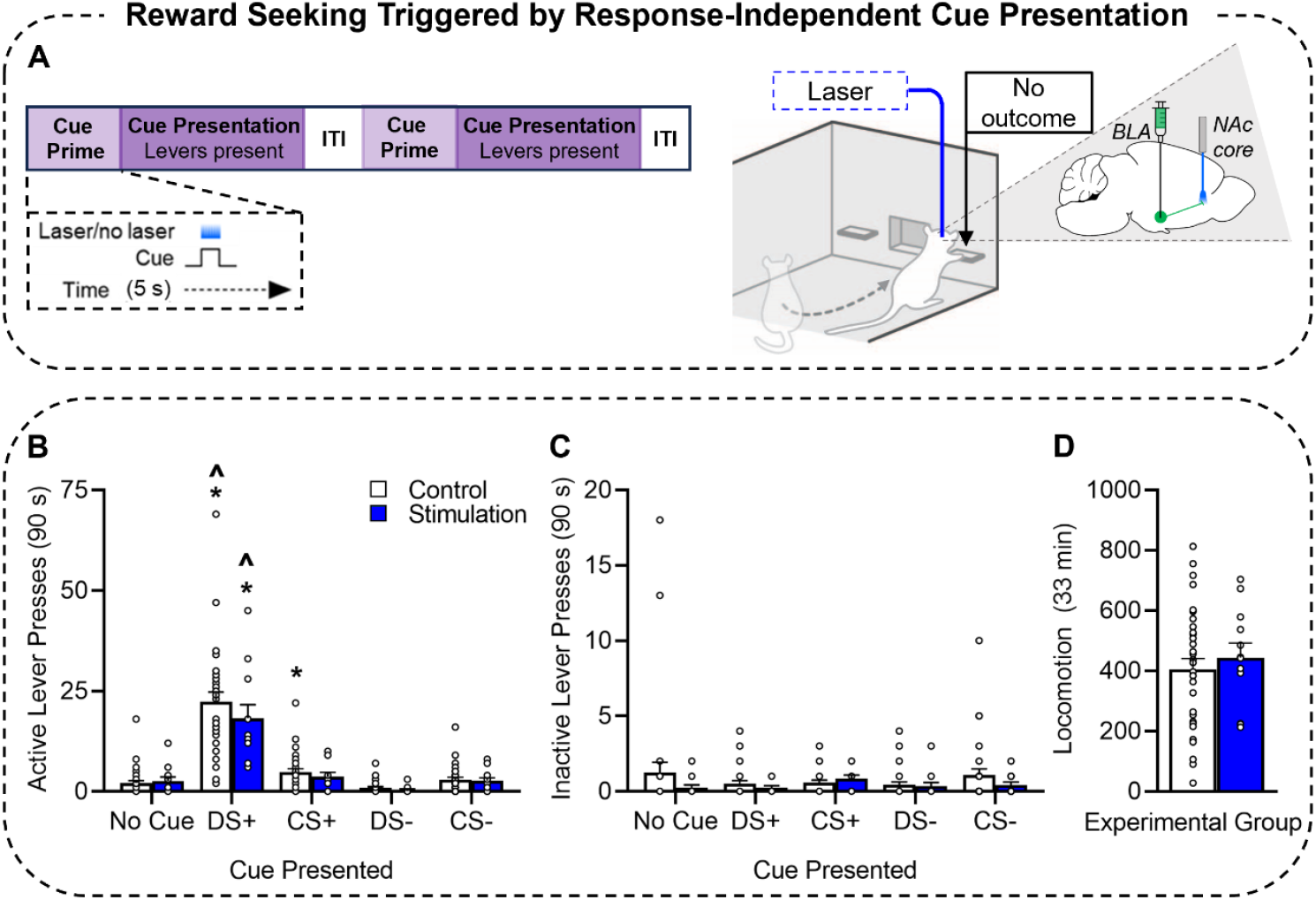
Effects of optogenetic stimulation of BLA neurons projecting to the NAc core on cue-evoked sucrose seeking. **(A)** Schematic of the cue-evoked sucrose seeking test where the DS+, CS+, DS-, CS-, and No Cue conditions were presented response-independently and paired with optogenetic stimulation. Lever pressing produced no cues and no sucrose. White bars represent control rats and blue bars represent rats receiving cue-paired optogenetic photostimulation. **(B)** Active lever presses, **(C)** inactive lever presses, and **(D)** locomotion. Data are average ± SEM. Herein, open circles in bars represent data from individual rats. DS+, discriminative stimulus previously signaling sucrose availability. DS-, discriminative stimulus previously signaling sucrose unavailability. CS+, conditioned stimulus paired with sucrose. CS-, conditioned stimulus never paired with sucrose. * vs. No Cue in the same group. ^ vs. CS+ in the same group. *n* = 32 Control, *n* = 12 Stimulation. (**A)** Adapted from Servonnet et al. (2023).

On one test, the 5-s priming period was paired with laser delivery (465 nm; 5 s, 10 mW, 5 ms pulses, 20 Hz); on the other test no laser was delivered. Test order was counterbalanced, and one discrimination retraining session was given between cue-evoked sucrose seeking tests to ensure that sucrose self-administration remained under the control of the DS+, as indicated by a DS+ discrimination ratio of ≥60% and self-administration of ≥6 sucrose reinforcers. In response to reward cues, BLA neurons fire in vivo at frequencies similar to the 20 Hz laser stimulation frequency we used here [51,52]. Thus, our optogenetic stimulation parameters produce reliable action potentials in ChR2-eYFP-expressing BLA neurons [8] and BLA→NAc core neurons [42] at physiologically relevant rates.

### Conditioned Reinforcing Properties of the Sucrose-Associated Cues

Here we tested the hypothesis that cue-paired BLA→NAc core neuron stimulation increases the instrumental pursuit of the DS+ and CS+ when the cues are presented contingent on lever pressing. Two days following the sucrose seeking tests, rats received 2 conditioned reinforcement tests during which lever pressing was now reinforced by presentation of the sucrose-associated cues, without sucrose reward. As illustrated in Fig 3A, conditioned reinforcement tests (33 min/session) began with a 10-s priming period (DS+, CS+, DS-, CS-, or a ‘No Cue’ condition; counterbalanced), followed by a 30-s period during which the prime was terminated, and the levers were inserted. During this 30-s period, active lever pressing produced a 5-s presentation of the condition that was presented during the priming period under an FR1 schedule. A VI 1.5-min ITI followed, during which no stimuli were presented and both levers were retracted. The ITI was followed by the next 10-s priming period and 30-s lever access period. This cycle repeated until rats could press to earn each cue condition 3 times. Active lever presses and the number of cues earned during lever access periods served as measures of the conditioned reinforcing properties of the cues.

**Fig 3.**
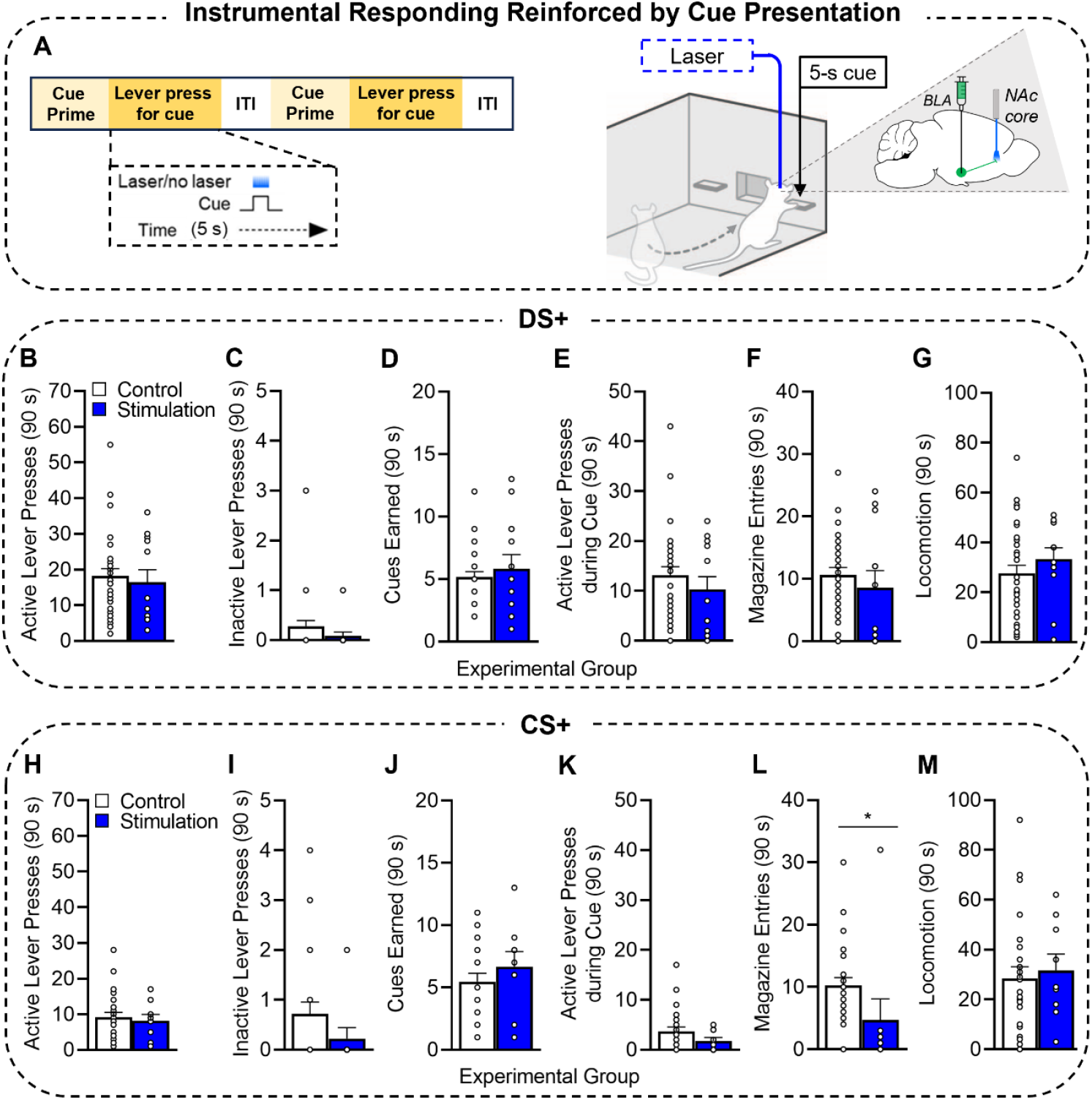
Effects of optogenetic stimulation of BLA neurons projecting to the NAc core on the conditioned reinforcing properties of a sucrose-associated DS+ and CS+. **(A)** Schematic of the conditioned reinforcement test where pressing the active lever was reinforced by cue presentation in the absence of sucrose. White bars represent control rats and blue bars represent rats receiving cue-paired optogenetic photostimulation. **(B)** Active lever presses to earn the DS+, **(C)** inactive lever presses, **(D)** DS+ presentations earned, **(E)** active lever presses during earned DS+ presentation, **(F)** magazine entries, and **(G)** locomotion during the 3 x 30-s trials where the DS+ could be earned. **(H)** Active lever presses to earn the CS+, **(I)** inactive lever presses, **(J)** CS+ presentations earned, **(K)** active lever presses during earned CS+ presentation, **(L)** magazine entries, and **(M)** locomotion during the 3 x 30-s trials where the CS+ could be earned. Data are average ± SEM. DS+, discriminative stimulus previously signaling sucrose availability. DS-, discriminative stimulus previously signaling sucrose unavailability. CS+, conditioned stimulus paired with sucrose. CS-, conditioned stimulus never paired with sucrose. ITI, intertrial interval. DS+: *n* = 32 Control, *n* = 12 Stimulation. CS+: *n* = 25 Control, *n* = 9 Stimulation. (**A)** Adapted from Servonnet et al. (2023).

On one test, each 5-s cue presentation earned was paired with laser (465 nm; 5 s, 10 mW, 5 ms pulses, 20 Hz), on the other test no laser was delivered. The tests proceeded in counterbalanced order, with an intervening discrimination retraining session.

### Experiment 2. Effects of Optogenetic Stimulation of BLA→PrL Neurons on Cue-Guided Sucrose Seeking

#### (DS+)-Evoked Increases in Sucrose Seeking

Here we tested the hypothesis that cue-paired BLA→PrL neuron stimulation enhances the ability of response-independent presentation of the DS+ to trigger increases in sucrose seeking. Fig 4A illustrates the test procedures, which proceeded as detailed under *‘(DS+)-Evoked Increases in Sucrose Seeking’* for Experiment 1, except that we photostimulated BLA→PrL neurons.

**Fig 4.**
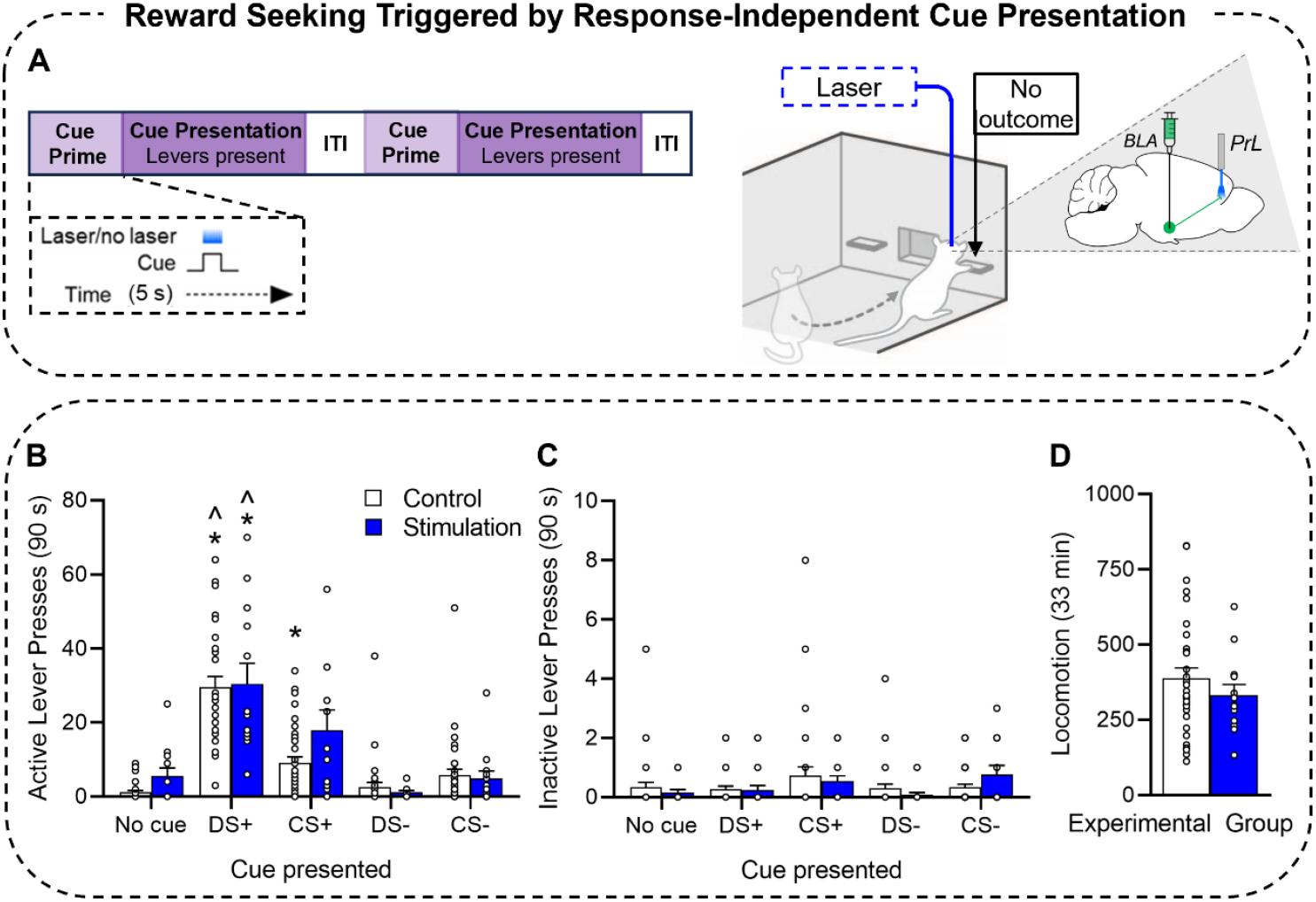
Effects of optogenetic stimulation of BLA neurons projecting to the PrL on cue-evoked sucrose seeking. **(A)** Schematic of the cue-evoked seeking test where all cue conditions were presented response-independently and paired with optogenetic stimulation. Lever pressing produced no cues and no sucrose. White bars represent control rats and blue bars represent rats receiving cue-paired optogenetic photostimulation. **(B)** Active lever presses, **(C)** inactive lever presses, and **(D)** locomotion. Data are average ± SEM. DS+, discriminative stimulus previously signaling sucrose availability. DS-, discriminative stimulus previously signaling sucrose unavailability. CS+, conditioned stimulus paired with sucrose. CS-, conditioned stimulus never paired with sucrose. * vs. No Cue in the same group. ^ vs. CS+ in the same group. *n* = 33 Control, *n* = 13 Stimulation. (**A)** Adapted from Servonnet et al. (2023).

#### Conditioned Reinforcing Properties of Reward Cues

Here we tested the hypothesis that cue-paired BLA→PrL neuron stimulation increases instrumental pursuit of the DS+ and CS+ when the cues are presented contingent on lever pressing. Fig 5A illustrates the test procedures, which were similar to those described under *‘Conditioned Reinforcing Properties of the Sucrose-Associated Cues’* for Experiment 1. Rats were initially tested under an FR3 schedule of cue reinforcement; however, this schedule produced very low levels of responding across cue types (data not shown). As a result, rats were retested under an FR1 schedule.

**Fig 5.**
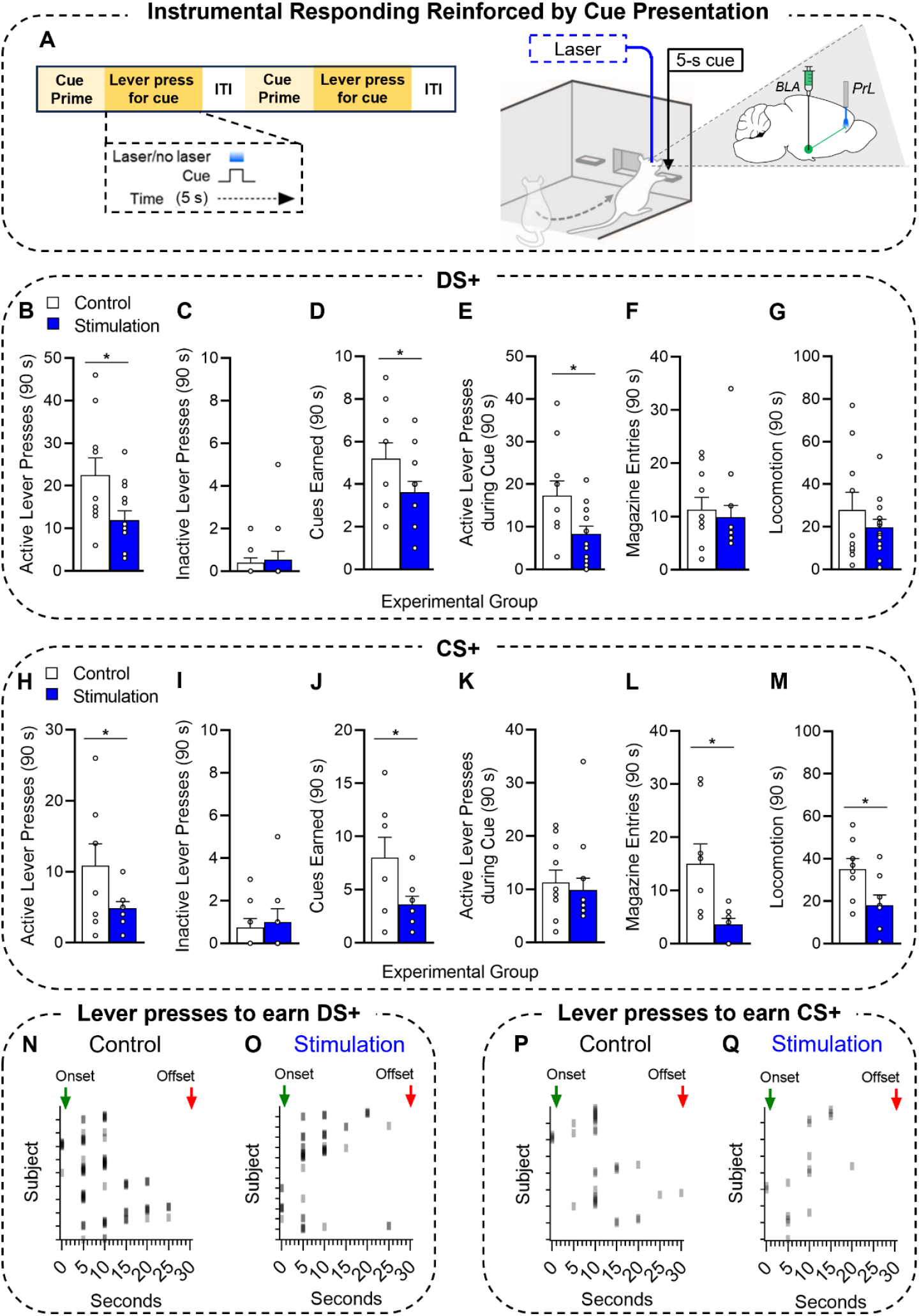
Effects of optogenetic stimulation of BLA neurons projecting to the PrL on the conditioned reinforcing properties of a sucrose-associated DS+ and CS+. **(A)** Schematic of the conditioned reinforcement test where pressing the active lever was reinforced by cue presentation, in the absence of sucrose. White bars represent control rats and blue bars represent rats receiving sucrose-paired optogenetic photostimulation. **(B)** Active lever presses to earn the DS+, **(C)** inactive lever presses, **(D)** DS+ presentations earned, **(E)** active lever presses during DS+ presentation, and **(F)** magazine entries and **(G)** locomotion during the 3 x 30-s trials where the DS+ could be earned. **(H)** Active lever presses to earn the CS+, **(I)** inactive lever presses, **(J)** CS+ presentations earned, **(K)** active lever presses during CS+ presentation, **(L)** magazine entries and **(M)** locomotion during the 3 x 30-s trials where the CS+ could be earned. Active lever presses to earn the DS+ in **(N)** control and **(O)** stimulation groups, and active lever presses to earn the CS+ in **(P)** control and **(Q)** stimulation groups. Data are average ± SEM. DS+, discriminative stimulus signaling sucrose availability. DS-, discriminative stimulus signaling sucrose unavailability. CS+, conditioned stimulus paired with sucrose. CS-, conditioned stimulus never paired with sucrose. ITI, intertrial interval. DS+: *n* = 10 Control, *n* = 13 Stimulation. CS+: *n* = 8 Control, *n* = 8 Stimulation. (**A)** Adapted from Servonnet et al. (2023).

#### Effects of Optogenetic Stimulation of BLA→PrL Neurons on the Reinforcing Properties of Sucrose

After determining the effects of BLA→PrL neuron stimulation on the conditioned reinforcing properties of the DS+ and CS+, we determined the extent to which stimulation influenced the reinforcing properties of the primary reinforcer, sucrose. Fig 6A illustrates the behavioural procedure. Rats received 2 discrimination training sessions as described above, except that now each session included 7, instead of 15, DS+/- trials to reduce risk of photostimulation-induced seizure activity. During one session, each sucrose delivery was paired with 5-s laser stimulation, and during the other session no laser was delivered. Session order was counterbalanced.

**Fig 6.**
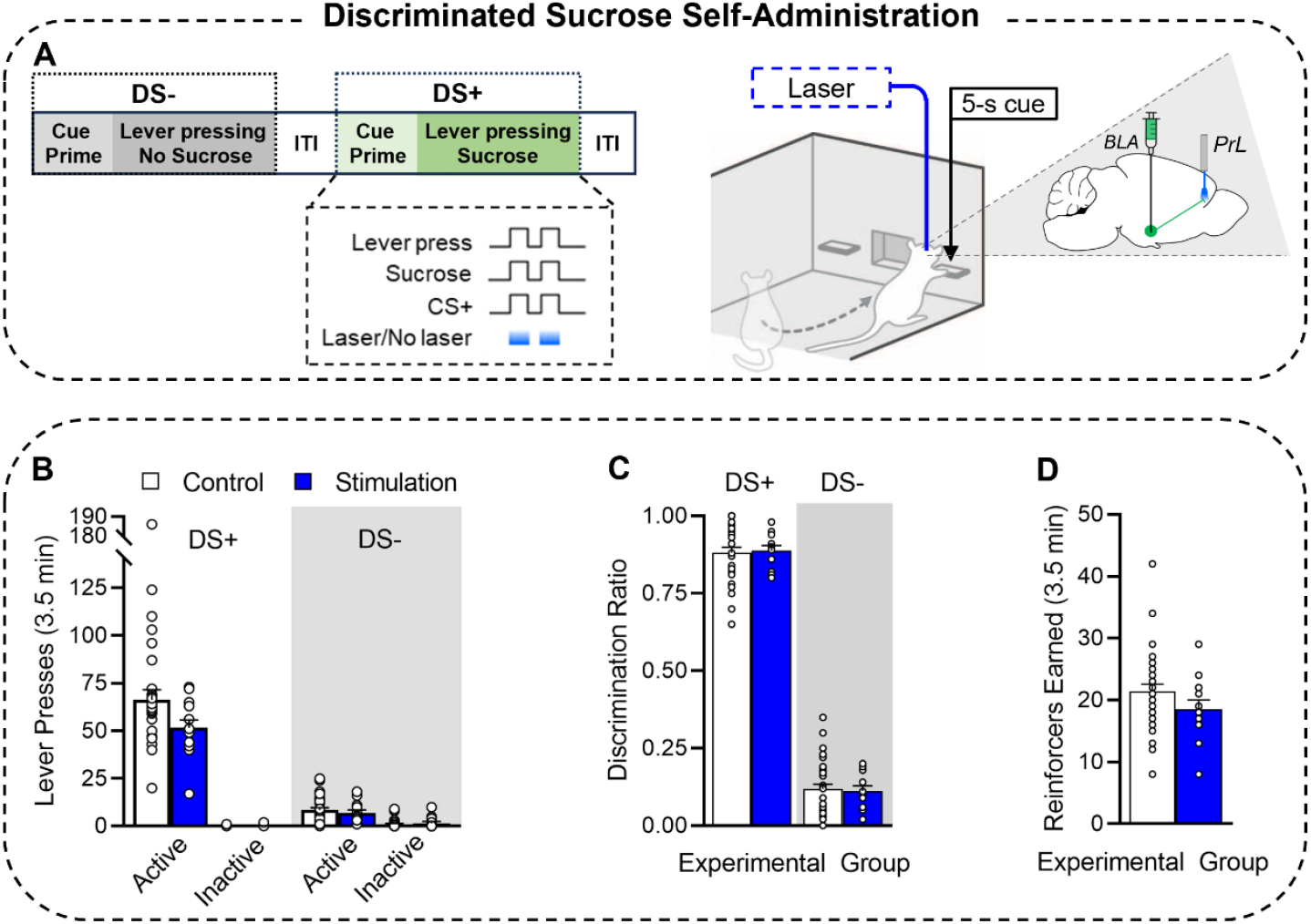
The effects of optogenetic stimulation of BLA neurons projecting to the PrL on sucrose self-administration under the control of discriminative stimuli. **(A)** Schematic of the discrimination training session during which pressing the active lever produced sucrose and a conditioned stimulus during DS+ trials and produced no sucrose and a CS-during DS-trials. Each sucrose delivery was paired with a 5-s photostimulation of BLA-to-PrL neurons. White represents control rats and blue represents rats receiving cue-paired optogenetic photostimulation. **(B)** Active and inactive lever presses during the DS+ and DS-, **(C)** discrimination ratios, and **(D)** sucrose reinforcers earned. *n* =33 Control, *n* = 13 Stimulation. Data are average ± SEM. DS+, discriminative stimulus signaling sucrose availability. DS-, discriminative stimulus signaling sucrose unavailability. ITI, intertrial interval. (**A)** Adapted from Servonnet et al. (2023).

### Experimental Design and Statistical Analyses

During conditioned reinforcement tests, we assessed whether pairing laser delivery with DS+ and CS+ presentations increases instrumental responding reinforced by the cues. Under these conditions, laser was delivered only upon cue delivery. To ensure a reliable test of our hypothesis, we included data only from rats that earned at least one presentation of each cue type. For example, if a rat did not earn a DS+ presentation, then DS+ related active lever presses, cues earned, and magazine entries were removed, while CS+ related data remained if at least one CS+ presentation was earned. Table 1 presents initial and final sample sizes in conditioned reinforcement tests across experiments. In addition, Table 2 shows initial and post-histology sample sizes across experiments.

**Table 1.** Sample sizes and attrition across conditioned reinforcement tests.

|  | DS<sup>+</sup> |  | CS<sup>+</sup> |  |
| --- | --- | --- | --- | --- |
|  | Initial | Final | Initial | Final |
| **Experiment 1** |  |  |  |  |
| Control | *n* = 32 | *n* = 32 | *n* = 32 | *n* = 25 |
| ChR2 | *n* = 12 | *n* = 12 | *n* = 12 | *n* = 9 |
| **Experiment 2** |  |  |  |  |
| Control | *n* = 33 | *n* = 33 | *n* = 33 | *n* = 31 |
| ChR2 | *n* = 13 | *n* = 13 | *n* = 13 | *n* = 8 |

**Table 2.**
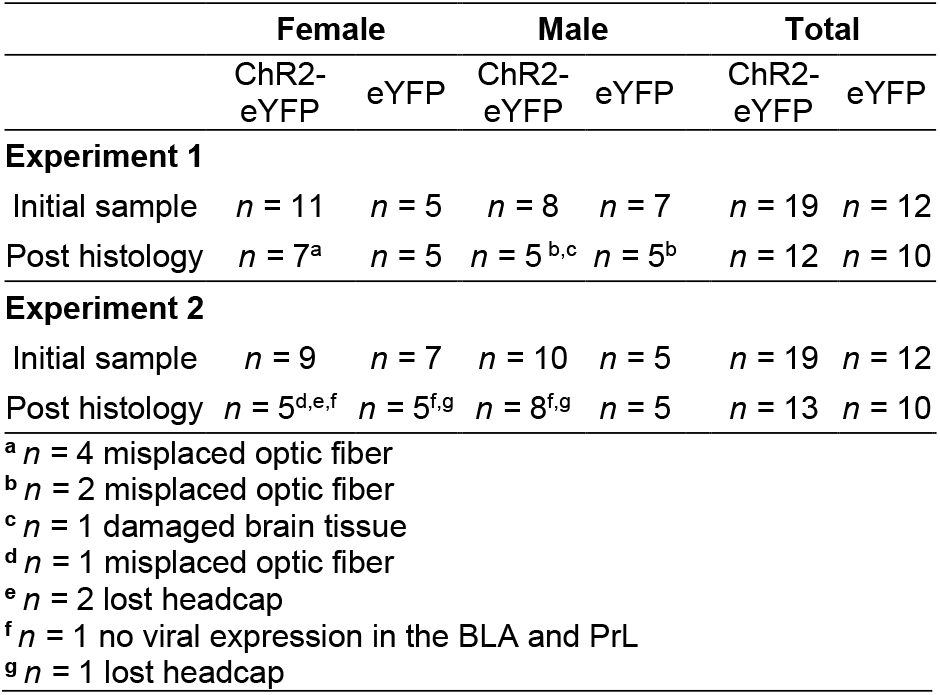
Initial and post-histology sample sizes.

|  | Female |  | Male |  | Total |  |
| --- | --- | --- | --- | --- | --- | --- |
|  | ChR2-eYFP | eYFP | ChR2-eYFP | eYFP | ChR2-eYFP | eYFP |
| **Experiment 1** |  |  |  |  |  |  |
| Initial sample | *n* = 11 | *n* = 5 | *n* = 8 | *n* = 7 | *n* = 19 | *n* = 12 |
| Post histology | *n* = 7<sup>a</sup> | *n* = 5 | *n* = 5<sup>b,c</sup> | *n* = 5<sup>b</sup> | *n* = 12 | *n* = 10 |
| **Experiment 2** |  |  |  |  |  |  |
| Initial sample | *n* = 9 | *n* = 7 | *n* = 10 | *n* = 5 | *n* = 19 | *n* = 12 |
| Post histology | *n* = 5<sup>d,e,f</sup> | *n* = 5<sup>f,g</sup> | *n* = 8<sup>f,g</sup> | *n* = 5 | *n* = 13 | *n* = 10 |
| <sup>a</sup> *n* = 4 misplaced optic fiber |  |  |  |  |  |  |
| <sup>b</sup> *n* = 2 misplaced optic fiber |  |  |  |  |  |  |
| <sup>c</sup> *n* = 1 damaged brain tissue |  |  |  |  |  |  |
| <sup>d</sup> *n* = 1 misplaced optic fiber |  |  |  |  |  |  |
| <sup>e</sup> *n* = 2 lost headcap |  |  |  |  |  |  |
| <sup>f</sup> *n* = 1 no viral expression in the BLA and PrL |  |  |  |  |  |  |
| <sup>g</sup> *n* = 1 lost headcap |  |  |  |  |  |  |

All experiments used a mixed design with Cue condition (DS+, CS+, DS-, CS-, or No Cue) as a within-subjects factor and biological sex (Female or Male) and treatment (Optogenetic stimulation or Control) as between-subjects factors. Unless specified in the results section, there were no significant sex differences in behaviour (Tables S2 and S3). We used Shapiro-Wilk tests to assess the normality of all data sets. We analysed normally distributed data using ANOVA or t-tests, and non-normally distributed data with Wilcoxon and Mann-Whitney tests. We used the Holm-Bonferroni correction for multiple comparisons. Statistical analyses were conducted with IBM SPSS (Version 26) and evaluated using a statistical significance level of p < 0.05. Graphs were created with GraphPad Prism (Version 8; La Jolla, CA).

Based on prior work [7,12–14,19,48,53,54] we predicted that compared to all other cue conditions, the DS+ would trigger greater peaks in lever pressing during cue-evoked sucrose seeking tests. Moreover, we predicted that stimulating BLA→PrL [34] or BLA→NAc core [37; c.f. 55, 42] projection neurons would increase the instrumental pursuit of the DS+ and CS+. Thus, a priori analyses were conducted on these comparisons.

## Results

### Sucrose self-administration under the control of discriminative stimuli

Sucrose self-administration and discrimination data were pooled across Experiments 1 & 2. Following the acquisition of sucrose self-administration behaviour (Fig S2), the DSs were presented in predictable order during discrimination training Sessions 1-9 and in unpredictable order from Session 10 onward. On Session 1 (Fig 1B), active lever presses peaked during DS-trials, where rats made 64% of their active lever presses (Fig 1B inset). However, by Session 9, responding peaked during DS+ trials (Fig 1C), with rats pressing on the active lever most (80%) during DS+ presentation, and responding little during DS-presentation (20% of total active lever presses; Fig 1C inset). On Session 10 the DSs were presented in unpredictable order for the first time. Nonetheless, rats continued to make most (78%) of their active lever presses during DS+ trials (Fig 1D inset) and DS discrimination remained strong until the last session (Fig 1E; 85% of total responding during DS+ trials).

Fig 1F further highlights the acquisition of discriminated responding, where the DS+ ratio increased (*t*_(44)_ = -24.39, *p* < 0.001) and the DS-ratio decreased (*t*_(44)_ = -24.47, *p* < 0.001) from the 1st to the last session. During the 1^st^ session, discrimination ratios were higher for the DS- vs. DS+ (*t*_(44)_ = -11.19, *p* < 0.001), indicating that rats initially pressed more on the active lever during DS-trials. However, by the last session, this pattern reversed, and rats now had higher DS+ vs. DS-ratios (*t*_(44)_ = -25.30, *p* < 0.001). To habituate the rats to tethering during optogenetic manipulations by, patch cords were attached to their optic fiber implants from session 14 onward. This had no effect on discrimination ratios. Thus, sucrose self-administration came under control of the DSs, whereby rats learned to self-administer sucrose during DS+ trials and to inhibit responding during DS-trials.

The only sex effect observed during sucrose self-administration was in the amount of sucrose self-administered (Fig 1G). From the 1st to the last session, males increased the number of sucrose reinforcers earned (*t*_(22)_ = -3.77, *p* = 0.001), whereas females showed no statistically significant change (*t*_(21)_ = -0.93, *p* = 0.36). Consequently, males earned more sucrose reinforcers than did females on the last session (*t*_(43)_ = -3.14, *p* = 0.003).

### Experiment 1. Effects of Optogenetic Stimulation of BLA→NAc core Neurons on Cue-Guided Sucrose Seeking

eYFP-expressing rats that received laser or no laser, and ChR2-eYFP-expressing rats that received no laser showed similar lever pressing behaviour. Thus, these groups were pooled into one control group in Experiment 1.

#### Cue-Evoked Increases in Sucrose Seeking

##### Responding in control rats

In control rats, cue type determined lever pressing behaviour. Even though lever pressing produced no outcome, rats pressed more on the active vs. inactive lever (Figs 2B-C) during presentations of the DS+ (*z* = -4.94, *p* < 0.001), CS+ (*z* = -4.19, *p* < 0.001), and CS^--^ (*z* = -2.99, *p* = 0.008). The rats pressed similarly on the two levers during the DS- (*z* = -1.27, *p* = 0.21) and No Cue (*z* = -1.84, *p* = 0.07) conditions. Compared to baseline levels of responding measured during the No Cue condition, rats pressed more on the active lever only during DS+ (*z* = -4.78, *p* < 0.001) and CS+ (*z* = -2.95, *p* = 0.009) presentations. The rats, however, responded significantly more to DS+ (Fig 2B) vs. CS+ (*z* = -4.81, *p* < 0.001), CS- (*z* = -4.84, *p* < 0.001), and DS- (*z* = -4.92, *p* < 0.001) presentations. Thus, a DS+ signaling sucrose availability triggered significant increases in sucrose-seeking behaviour in the absence of the primary reward and was more effective compared to a CS^+^ signaling sucrose delivery.

Across Experiments 1 and 2, neither Pavlovian conditioned approach phenotype (Tables S4 and S5) nor breakpoint for sucrose (Table S6) significantly correlated with responding during the cue-induced sucrose seeking test. Thus, neither incentive motivation for sucrose nor a sign- vs. goal-tracking phenotype significantly predicted cue-evoked sucrose seeking behaviour.

##### Effects of optogenetic stimulation of BLA→NAc core projection neurons

Cue type also determined responding in rats receiving cue-paired photostimulation of BLA→NAc core neurons (Figs 2B-C). The rats pressed more on the active vs. inactive lever (Figs 2B-C) during the DS+ (*z* = - 3.07, *p* = 0.005) and CS- (*z* = -2.58, *p* = 0.04). In contrast, rats showed no lever discrimination during CS+ (*z* = -2.10, *p* = 0.11), DS- (*z* = -0.28, *p* = 0.78) or No Cue (*z* = -2.25, *p* = 0.07) conditions.

Compared to controls, cue-paired photostimulation of BLA→NAc core neurons had no effect on responding to the DS+ or CS+ (Fig 2B: DS+: *z* =-1.04, *p* = 0.30; CS+: *z* =-0.84, *p* = 0.41). Photostimulation also did not significantly affect locomotion (Fig 2D) during testing relative to control (*t*_*(42)*_ = - 0.60, *p* = 0.56). Thus, optogenetic stimulation of BLA→NAc core projection neurons did not influence cue-evoked increases in sucrose-seeking responses.

#### The Conditioned Reinforcing Properties of the DS+ and CS+

##### Responding in control rats

Control rats pressed more on the active vs. inactive lever across all cue conditions (Figs S3B-C; DS+: *z* = -4.94, *p* < 0.001; CS+: *z* = -4.39, *p* < 0.001; DS-: *z* = -2.93, *p* = 0.003; CS-: *z* = -3.85, *p* < 0.001; No cue: *z* = -3.73, *p* < 0.001). However, relative to baseline levels of responding measured during the No Cue condition, rats pressed more on the active lever to earn the DS+ (*z* = -4.94, *p* < 0.001) and CS+ (*z* = -2.75, *p* = 0.009), but not the DS- (*z* = -0.43, *p =* 0.67) or CS- (*z* = -0.78, *p* = 0.43). Thus, only the DS+ and CS+ acquired significant conditioned reinforcing value. Importantly, rats responded more to earn the DS+ vs. the CS+ (*z* = -4.62, *p* < 0.001). However, they earned similar numbers of DS+ and CS+ presentations (Fig S3D; *z* = 1.29, *p* = 0.20). The higher rate of responding under the DS+ condition reflects increased lever pressing *during* cue presentation—even though this produced no outcome—(Fig S3E). During earned DS+ presentations, the rats increased their rate of active lever pressing compared to both the No Cue (*z* = -4.86, *p* < 0.001) and CS+ (*z* = - 4.73, *p* < 0.001) conditions. Active lever pressing during earned CS+ presentation was similar to that during the No cue condition (*z* = -0.22, *p* = 0.83). Thus, when instrumental responding was reinforced by a sucrose-associated DS+ or CS+, the DS+ supported the highest rates of responding, including during its presentation, suggesting more robust conditioned reinforcing properties.

Pavlovian conditioned approach phenotype (Tables S4 and S5) nor breakpoint for sucrose (Table S6) significantly correlated with responding during conditioned reinforcement tests across experiments. Thus, neither a sign- or goal-tracking phenotype nor incentive motivation for sucrose were significant predictors of the instrumental pursuit of sucrose-associated cues.

##### Effects of optogenetic stimulation of BLA→NAc core projection neurons

Just like control rats, rats receiving cue-paired photostimulation of BLA→NAc core neurons also showed higher rates of responding on the active vs. inactive lever across cue conditions (Figs 3B-C, DS+: *z* = -4.94, *p* < 0.001; Figs 3H-I, CS+: *z* = -4.39, *p* < 0.00. Data not shown; DS-: *z* = -2.93, *p* = 0.003; No Cue: *z* = -3.73, *p* < 0.001; CS- : *z* = -3.85, *p* < 0.001).

Compared to controls, photostimulation had no effect on instrumental responding for the sucrose-associated DS+ and CS+. Indeed, there were no effects of photostimulation on total active lever presses (Fig 3B, DS+: *z* = -0.46, *p* = 0.65; Fig 3H, CS+: *z* = -0.16, *p* = 0.88), number of cues earned (Fig 3D, DS+: *z* = -0.63, *p* = 0.54; Fig 3J, CS+: *z* = - 0.63, *p* = 0.54), active lever presses during cue presentation (Fig 3E, DS+: *z* = -0.81, *p* = 0.43; Fig 3K, CS+: *z* = -1.10, *p* = 0.30), or magazine entries during the 3 x 30-s trials where DS+ presentation could be earned (Fig 3F, *z* = -1.12, *p* = 0.27). However, photostimulation reduced magazine entries during the 3 x 30-s trials where the CS+ could be earned (Fig 3L; *z* = -3.15, *p* = 0.001). Finally, there was no effect of photostimulation on locomotion during the during the 3 x 30- s trials where DS+ (Fig 3G, *z* = -1.17, *p* = 0.25) or CS+ (Fig 3M, *z* = -0.60, *p* = 0.62) presentations could be earned. Thus, photostimulation of BLA→NAc core projection neurons had no effect on the reinforcing properties of a sucrose-associated DS+ or CS+, but it suppressed conditioned approach behaviour during self-administered CS+ presentations.

### Experiment 2. Effects of Optogenetic Stimulation of BLA→PrL Neurons on Cue-Guided Sucrose Seeking

#### Cue-Evoked Increases in Sucrose Seeking

During cue-evoked seeking tests, lever pressing was similar across eYFP-expressing rats receiving laser or no laser and ChR2-eYFP-expressing rats receiving no laser. These groups were therefore pooled into a single control group.

#### Responding in control rats

Across cue conditions, control rats pressed more on the active lever vs. inactive lever (Figs 4B-C; DS+: *z* = -5.01, *p* < 0.001; CS+: *z* = -4.27, *p* < 0.001; DS-: *z* = -2.61, *p* = 0.02; CS-: *z* = -4.37, *p* < 0.001; No Cue: *z* = -2.12, *p* = 0.03), even though lever pressing produced no outcome. Cue condition also determined responding on the active lever (Fig 4B). Relative to the No Cue condition, rats responded more during DS+^+^ (*z* = -5.01, *p* < 0.001), CS+ (*z* = -4.19, *p* < 0.001), and CS- (*z* = - 4.16, *p* < 0.001) presentations. They also responded most during DS+ presentation (vs. CS+: *z* = -4.75, *p* < 0.001; vs. CS-: *z* = - 4.95, *p* < 0.001, vs. DS-: *z* = -5.03, *p* < 0.001). Thus, as in Experiment 1 (Fig 2), the DS+ was the most effective cue type in triggering peaks in sucrose-seeking behaviour.

#### Effects of optogenetic stimulation of BLA→PrL projection neurons

Rats receiving cue-paired photostimulation responded more on the active vs. inactive lever (Figs 4B-C) during DS+ (*z* = -3.18, *p* = 0.004) and CS+ (*z* = -2.94, *p* = 0.007) presentation. There was no lever discrimination during DS- (*z* = -2.23, *p* = 0.08), CS- (*z* = -2.37, *p* = 0.053) or No Cue (*z* = -2.21, *p* = 0.08) presentation.

Compared to control, optogenetic stimulation had no effect on active lever pressing in response to DS+ or CS+ presentation (Fig 4B; DS+: *z* = -0.23, *p* = 0.082; CS+: *z* = - 0.92, *p* = 0.36). Photostimulation also did not significantly affect locomotion (Fig 4D) relative to control (*z* = -1.01, *p* = 0.31). Thus, optogenetic activation of BLA→PrL projection neurons did not significantly influence cue-evoked increases in sucrose-seeking responses.

#### The Conditioned Reinforcing Properties of the DS+ and CS+

During tests of instrumental responding for conditioned reinforcement, all ChR2-expressing rats received laser and the control group consisted of eYFP-expressing rats receiving laser or no laser.

#### Responding in control rats

Control rats pressed more on the active vs. inactive lever to earn DS+ (*z* = -2.81, *p* = 0.01) and CS+ (*z* = -2.52, *p* = 0.02) presentations (Figs S4B-C). There was no lever discrimination under the other cue conditions (No Cue: *z* = 0.75, *p* = 0.08; DS-: *z* = -0.21, *p* = 0.83; CS-: -2.32, *p* = 0.06). Cue condition also determined responding on the active lever (Fig S4B). Rats responded more to earn the DS+ relative to both the No Cue (*z* = -2.81, *p* = 0.01) and CS+ conditions (*z* = -2.30, *p* = 0.02). In contrast, responding for the CS+ (*z* = -1.60, *p* = 0.16), CS- (*z* = -1.89, *p* = 0.24) and DS- (*z* = -0.67, *p =* 0.50) was similar to that during the No Cue condition. However, rats earned similar numbers of DS+ and CS+ presentations (Fig S4D; *z* = -0.38, *p* = 0.72). Rats responded more on the active lever during earned DS+ presentations than they did during earned CS+ presentations (*z* = -2.80, *p* = 0.005) or during the No Cue condition (Fig S4E; *z* = -2.81, *p* = 0.02). However, they did not press more during earned CS^+^ presentations than they did during the No Cue condition (*z* = -0.84, *p* = 0.40). Thus, as in Experiment 1, the DS+ and CS+ gained conditioned reinforcing properties such that both cue types reinforced instrumental responding in the absence of the primary sucrose reward. However, the DS^+^ supported higher rates of instrumental responding, particularly during its presentation.

#### Effects of optogenetic stimulation of BLA→PrL projection neurons

Rats receiving cue-paired optogenetic stimulation of BLA→PrL projection neurons also responded more on the active vs. inactive lever for the DS+ (*z* = -3.11, *p* = 0.009; Fig 5B vs. C). However, there was no lever discrimination under the other cue conditions (Fig 5H vs. I CS+: *z* = -2.07, *p* = 0.12. Data not shown: DS-: *z* = -1.48, *p* = 0.14; CS-: *z* = -2.31, *p* = 0.08; No Cue: *z* = -0.96, *p* = 0.34).

Compared to the controls, rats receiving photostimulation responded less on the active lever to earn the DS+ (Fig 5B; *t*_(21)_ = 2.43, *p* = 0.01), self-administered fewer DS+ presentations (Fig 5D: *t*_(21)_ = 1.83, *p* = 0.04), and responded less on the active lever during earned DS+ presentations (Fig 5E: *t*_(21)_ = 2.43, *p* = 0.01). During the 3 x 30-s trials where the DS+ could be earned, photostimulation had no significant effect on magazine entries (Fig 5F; *z* = -0.66, *p* = 0.52) or locomotor behaviour (Fig 5G: *t*_(12.61)_ = 0.96, *p* = 0.35). Photostimulation of BLA→PrL projection neurons had similar effects on responding for the CS+. Rats receiving photostimulation responded less on the active lever to earn the CS+ (Fig 5H; *t*_(21)_ = 1.88, *p* = 0.048) and self-administered fewer CS+ presentations (Fig 5J; *t*_(21)_ = 2.11, *p* = 0.03). However, photostimulation did not influence active lever presses during earned CS+ presentation (Fig 5K; *z* = -0.76, *p* = 0.25). In contrast to what was seen with the DS+, during the 3 x 30-s trials where CS+ presentations could be earned, photostimulation significantly reduced both magazine entries (Fig 5L; *t*_(21)_ = 2.90, *p* = 0.02) and locomotor behaviour (Fig 5M: *t*_(14)_ = 2.49, *p* = 0.03).

Figs 5N-Q further highlight the suppressive effects of BLA→PrL neuron photostimulation on responding for conditioned reinforcement. Figs 5N & O illustrate responding to the first DS+ in respectively, individual control rats and rats receiving photostimulation. Similarly, Figs 5P & Q show responding to the first CS+ in, respectively, individual control rats and rats receiving photostimulation. When each earned DS+ or CS+ was paired with optogenetic stimulation of BLA→PrL neurons, rats responded less to self-administer these cues.

Thus, photostimulation of BLA→PrL projection neurons significantly reduced the reinforcing properties of both the DS+ and CS+. Stimulation also had cue-specific effects, reducing seeking responses during earned DS+ but not CS+ presentation, and reducing conditioned approach behaviour to the site of reward delivery during CS+ but not DS+ presentation.

#### Effects of optogenetic stimulation of BLA→PrL Projection Neurons on the Reinforcing Properties of Sucrose

To determine the extent to which photostimulation of BLA→PrL neurons influences primary reinforcement, rats received a sucrose self-administration session during which each sucrose delivery was paired with photostimulation. Rats that received photostimulation did not significantly differ from controls in their rates of active lever pressing during DS+ or DS-trials (Fig 6B; DS+: *z* = -1.50, *p* = 0.13; DS-: *z* = -0.38, *p* = 0.70). Photostimulation also had no effect on discrimination ratios (Fig 6C; DS+: *z* = -0.32, *p* = 0.75; DS-: *z* = -0.32, *p* = 0.75), or the number of sucrose reinforcers earned (Fig 6D; *t*_(44)_ = -0.71, *p* > 0.48). Thus, photostimulating BLA→PrL projection neurons had no effect on the primary reinforcing properties of sucrose or the ability of the rats to use discriminative stimuli to guide their sucrose self-administration behaviour.

### Histology and c-Fos expression in the PrL induced by stimulation of ChR2-eYFP-expressing BLA terminals

Figs 7A-B show BLA cell bodies expressing, respectively, eYFP and ChR2-eYFP. Figs 7C-D show, respectively, eYFP- and ChR2-eYFP-expressing BLA terminals in the NAc core and outlines of the optic fibre implants. Figs 7F-G show, respectively, eYFP- and ChR2-eYFP-expressing BLA terminals in the PrL and outlines of the optic fibre implants. Figs E and H show, respectively, the placement of optic fibers in the NAc and PrL.Figs 7I-J show c-Fos protein expression in the PrL of an eYFP-expressing rat in the control hemisphere and the hemisphere with an optic fiber for photostimulation. Figs 7K-L show the same for a ChR2-eYFP-expressing rat. Fig 7M shows the ratio of c-Fos+ cells in the PrL hemisphere receiving optogenetic stimulation relative to the control hemisphere, in eYFP and ChR2-eYFP rats. This c-Fos ratio was higher in ChR2-eYFP-expressing rats receiving stimulation compared to eYFP-expressing rats (*z* = -2.80, *p* = 0.002). Thus, stimulating ChR2-eYFP expressing BLA terminals in the PrL increased the number of c-Fos immunoreactive cells in the PrL cortex.

**Fig 7.**
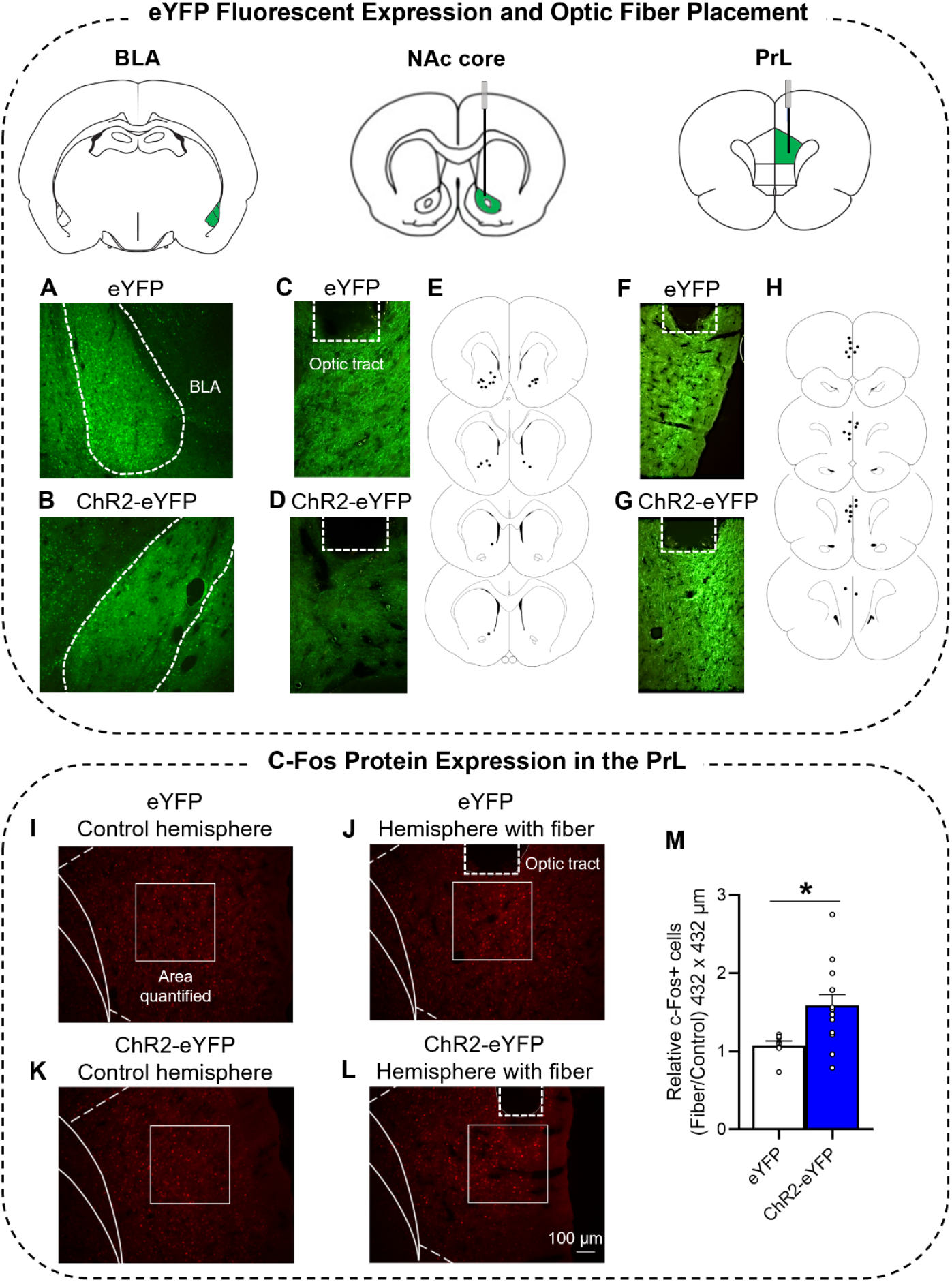
Confirmation of eYFP expression and optic fiber tracts in target brain regions, and induction of c-Fos protein expression in the PrL by optogenetic stimulation of BLA→PrL projection neurons. Representative images of **(A)** eYFP- and **(B)** ChR2-eYFP-expressing cell bodies in the BLA, **(C)** eYFP and **(D)** ChR2-eYFP terminal expression and **(E)** optic fiber placement in the NAc core, and **(F)** eYFP and **(G)** ChR2-eYFP terminal expression and **(H)** optic fiber placement in the PrL. Representative images of c-Fos expression in the PrL of an eYFP-expressing rat in the **(I)** control hemisphere containing no optic fiber and the **(J)** hemisphere receiving optogenetic stimulation via an optic fiber, and from a ChR2-eYFP-expressing rat in the **(K)** hemisphere containing no optic fiber and the **(L)** hemisphere receiving optogenetic stimulation via an optic fiber. **(M)** Average (± SEM) ratio of c-Fos positive cells in the optic fiber-containing hemisphere vs. contralateral hemisphere in control rats (white bar) and ChR2-expressing rats (blue bar). eYFP; *n* = 7 rats with AAV2/7-CamKIIa-EYFP virus, ChR2; *n* = 11 rats with AAV2/7-CamKIIa-hChR2(H134R)-EYFP virus.

## Discussion

We investigated the roles of BLA neurons projecting to the NAc core and the PrL in two forms of cue-motivated behaviour: *(i)* increases in sucrose-seeking behaviour triggered by response-independent DS and CS presentation, and *(ii)* reinforcement of instrumental responding by response-dependent DS and CS presentation. Rats learned that a DS^+^ signalled when instrumental responding would be reinforced with sucrose, and that a CS^+^ predicted sucrose reward delivery after a response. When the DS+ and CS+ were later presented response-contingently, they were equally effective as conditioned reinforcers of sucrose-seeking behaviour, but when presented non-contingently, the DS+ triggered greater increases in reward seeking than did the CS+. Optogenetic stimulation revealed projection-specific functions: BLA→NAc core stimulation had no detectable effect on either cue motivated behaviour, whereas BLA→PrL stimulation selectively suppressed the conditioned reinforcing effects of the DS+ and CS+, without altering sucrose self-administration behaviour. These results suggest that BLA→PrL activity reduces the incentive value of reward-associated cues, making them less desirable and thereby constraining cue-motivated reward seeking.

### The ability of a DS vs. a CS to act as a Pavlovian conditioned excitor of reward-seeking behaviour

When presented unexpectedly and independently of responding, the DS+ robustly increased sucrose seeking, whereas the CS+ did not. This extends prior findings that DSs increase reward seeking across reward types [7,12,38,53,56,13–15,17–21]. Our findings also support the idea that cues signalling when seeking actions will be reinforced (i.e., DSs) acquire stronger incentive motivational properties than do cues paired with reward delivery (i.e., CSs). Under this framework, DSs serve as powerful Pavlovian conditioned excitors that promote reward seeking in the absence of primary reinforcement [57]. Because all animals, including humans, frequently encounter response-independent DSs in daily life, our findings have broad implications for understanding cue-guided reward-seeking behaviour.

Beyond triggering reward-seeking behaviour, reward-associated cues can also act as conditioned reinforcers of behaviour when they are presented response-contingently, sustaining instrumental responding when reward is not immediately available. Here, both the DS+ and CS+ reinforced responding (whereas a DS^-^ and CS^-^ did not) and rats earned comparable numbers of each cue presentation, indicating that the two cue types acquired similar conditioned incentive value through associative learning [9,58]. These findings replicate the well-established conditioned reinforcing effects of CSs and extend evidence that DSs can also acquire such effects [7,12,13,20,59]. While it remains to be determined whether DSs serve as conditioned reinforcers in humans, everyday scenarios— such as repeatedly checking the time before a meal— suggest that they may. Thus, our findings reveal a dissociation: whereas both reward-associated DSs+ and CSs+ reinforce instrumental responding, only DSs+ effectively prompt peaks in reward-seeking behaviour, reflecting overlapping yet distinct motivational processes.

### Effects of BLA→NAc core stimulation on DS+ and CS+

Cue-paired optogenetic stimulation of BLA→NAc core neurons did not alter (DS+)-evoked increases in reward seeking or instrumental responding reinforced by the DS+ or CS+. This contrast with reports that inactivation of BLA→NAc core projections reduces CS-reinforced responding [35,36]. Because our stimulation parameters reliably evoke action potentials in BLA→NAc core neurons [42], the absence of behavioural effects is unlikely due to inadequate activation of these neurons. Instead, subregion specificity within the NAc may be involved. Prior studies did not distinguish between BLA inputs to the NAc core vs. shell (e.g., [35]; see Fig 2A in [36]), whereas we selectively targeted the core. Our findings are consistent with recent evidence that BLA→NAc core stimulation does not enhance the ability of a CS+ to support learning of a new instrumental action [42]—another measure of conditioned reinforcement. Together, these results suggest that increasing activity in the BLA→NAc core pathway is not sufficient to enhance the ability of a DS+ to trigger increases in reward seeking, or the ability of a DS+ or CS+ to act as a conditioned reinforcer of reward seeking. Thus, the BLA projections involved likely terminate outside the NAc core.

### BLA→PrL stimulation does not influence increases in reward seeking behaviour evoked by a DS+

This null effect cannot be attributed to insufficient circuit activation, as the same stimulation parameters increased c-Fos expression in the PrL of ChR2-eYFP-expressing rats and suppressed instrumental responding reinforced by the DS+ and CS+ (see Fig 5). These findings do not support our hypothesis that enhancing BLA→PrL activity amplifies the effects of a DS+ as a Pavlovian conditioned excitor of reward seeking, and align with prior evidence that PrL activity is not required for DS+ control of cocaine taking and seeking behaviour [18,60]. Instead, the PrL may preferentially contribute to inhibitory conditioned control over behaviour. Indeed, inactivating the PrL disinhibits reward-seeking behaviour during periods where it is normally low, such as during presentation of a DS^-^, neutral stimuli, or intervals with no cues [27,39,40]. Thus, when cues are encountered independently of seeking actions, the BLA→PrL pathway might preferentially contribute under conditions of conditioned inhibition rather than conditioned excitation of reward seeking.

### BLA→PrL stimulation suppresses the instrumental pursuit of reward-associated DS+ and CS+

This can involve several mechanisms. First, photostimulation could non-specifically reduce motor function. However, this is unlikely because photostimulation decreased lever pressing reinforced by the DS+ without influencing general locomotion. While reductions in both locomotion and (CS+)-evoked magazine entries accompanied photostimulation-induced decreases in lever pressing for the CS+, similar photostimulation parameters produced no locomotor effects during cue-evoked sucrose seeking tests. This suggests that the reduced locomotion was a consequence rather than a cause of diminished CS+ responding. Second, photostimulation could have decreased sucrose value or produced aversive effects. This is also unlikely, as BLA→PrL stimulation did not significantly alter sucrose self-administration behaviour. Finally, photostimulation could have simply disrupted the normal flow of information within BLA→PrL neurons that otherwise occurs as animals engage in an instrumental responding task. We do not believe this to be the case, because photostimulation had no effect on cue-evoked increases in sucrose seeking or on sucrose self-administration behaviour. Instead, our findings suggest that BLA→PrL activity constrains the instrumental pursuit of sucrose-associated cues, likely by modulating neural representations of DS+ and CS+ and/or cue-reward associations [61].

Our results appear inconsistent with studies showing that inhibition of the BLA→PrL pathway reduces the ability of drug-paired CSs to reinforce reward seeking behaviour [33,34]. Several methodological differences could be involved, including reward type (sucrose here vs. cocaine in [33, 34], the BLA subregion targeted (posterior here vs. anterior in [33], and the presence or absence of extinction training, which can influence the neural mechanisms supporting conditioned reinforcement [62–65]. Another important distinction is that we stimulated, rather than inhibited BLA→PrL projections. Stimulation and inhibition of the same projection do not necessarily produce opposite outcomes, as behavioural effects depend on how downstream neurons and circuits are modulated. Using a CamKIIa promoter to drive opsin expression, our approach targeted putative glutamatergic BLA→PrL projection neurons [66], some of which synapse onto GABAergic interneurons that inhibit pyramidal cell activity [67,68]. Thus, our photostimulation conditions might have also activated inhibitory neurons in the PrL, potentially producing a net suppression of PrL outputs and a consequent dampening of DS+ and CS+ conditioned reinforcing effects. Alternatively, stimulation of BLA→PrL neurons in our study may have biased PrL output toward downstream targets that inhibit cue-driven behavior, such as the paraventricular thalamus [69,70]. Future studies can examine these hypotheses.

## Conclusions

Our findings identify circuit-specific contributions of BLA outputs in mediating the motivational properties of reward-predictive cues. BLA→NAc core stimulation did not influence (DS+)- or (CS+)-guided sucrose seeking. In contrast, BLA→PrL stimulation selectively suppressed instrumental responding reinforced by sucrose-associated DSs and CSs, without affecting self-administration of the primary reward. This could involve enhanced recruitment of local inhibitory interneurons within the PrL and/or recruitment of PrL-dependent projection neurons that might dampen cue responding, such as PrL→paraventricular thalamus projections. Together, these results reveal a novel inhibitory role for BLA→PrL projections in constraining the conditioned reinforcing properties of reward-predictive cues, reducing their ability to maintain seeking responses. To the extent that these mechanisms translate to humans, our findings provide evidence-based insight into how maladaptive cue-guided pursuit of reward, as seen in addiction, binge eating, and gambling disorders, could be targeted at the circuit level.

## Acknowledgements

This work was supported by grants to A.-N.S. from the National Science and Engineering Research Council of Canada (NSERC; Grant 355923) and the Courtois Fund. M.R.L. was supported by a postdoctoral fellowship from NSERC. ChatGPT was used to improve the clarity of the manuscript text.

