## Supplemental Materials for "Basolateral Amygdala Projections to the Prelimbic Cortex Suppress the Motivational Value of Reward-Associated Cues"

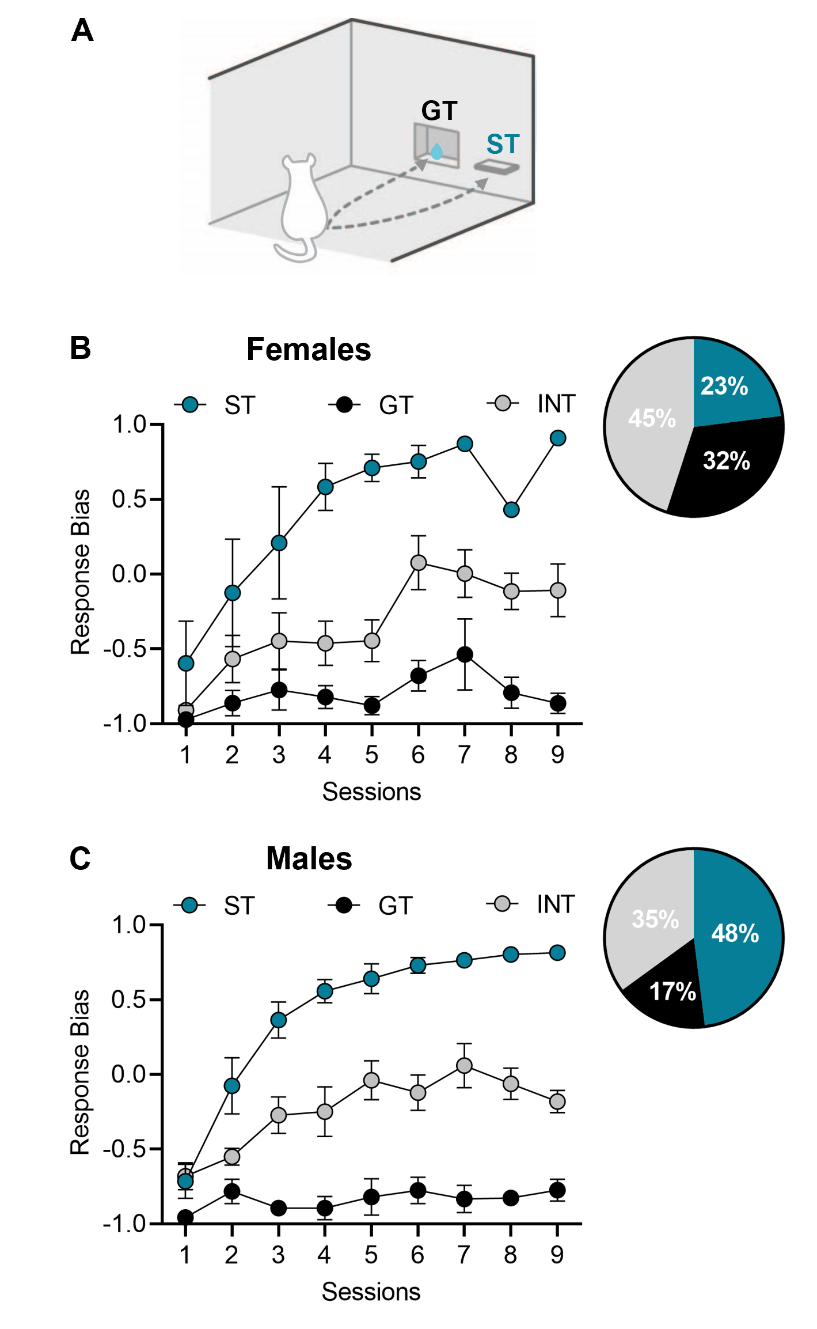

**Supplementary Figure 1. Acquisition of Pavlovian conditioned approach responding collapsed across Experiments 1 and 2. (A)** Schematic of the autoshaping procedure where rats learned that a lever conditioned stimulus predicts liquid sucrose. Average (± SEM) response bias for sign-tracking (ST; blue circles), goal-tracking (GT; black circles), and intermediate (INT; grey circles) Pavlovian conditioned approach phenotypes in **(B)** females and **(C)** males. Pie chart insets represent percentage of rats with each phenotype. Rats in Experiment 1 received 6 sessions. Rats in Experiment 2 received 9 sessions. Schematic in **(A)** adapted from Servonnet et al. (2023).

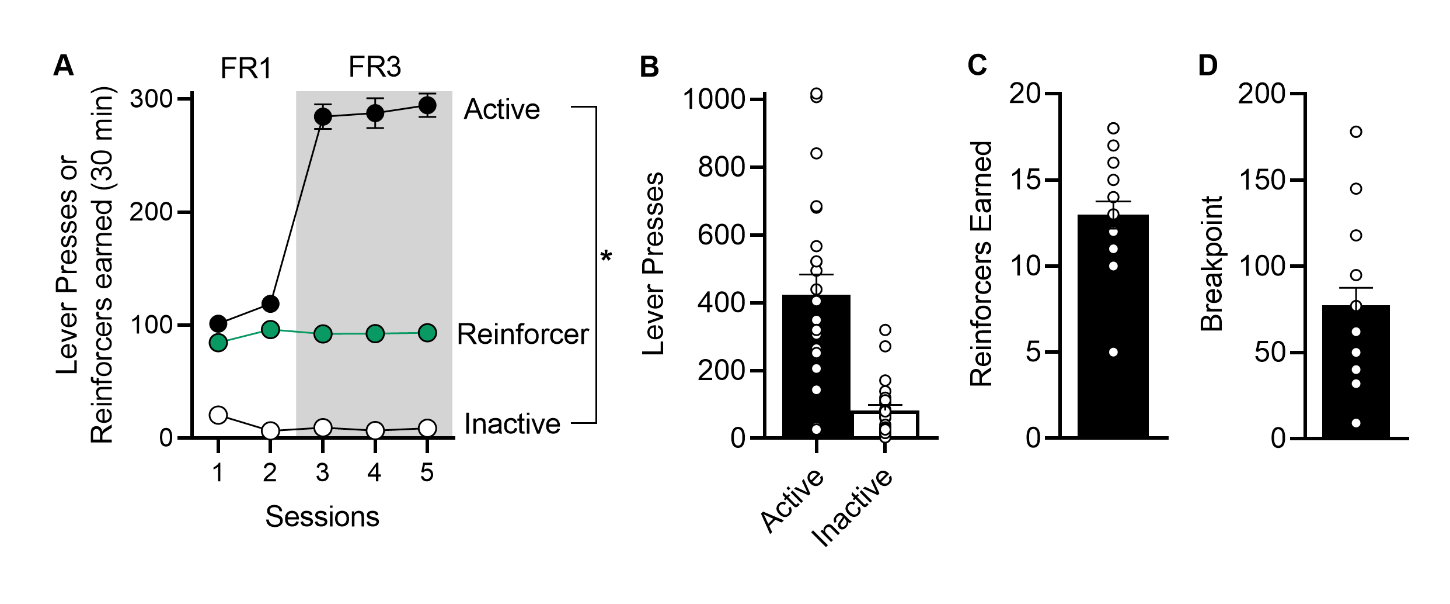

**Supplementary Figure 2.** **Acquisition of sucrose self-administration.** Rats pressed more on the active versus inactive lever during sucrose self-administration sessions (Session 1: (*t*_(44)_ = -20.25, *p* < 0.001; Session 2: *z* = -5.84, *p* < 0.001; Session 3: *z* = -5.84, *p* < 0.001; Session 4: *z* = -5.84, *p* < 0.001; Session 5: *z* = -4.20, *p* < 0.001). Inactive lever presses decreased across sessions (Session 1 vs. Last session: *z* = -4.31, *p <* 0.001). Rats earned ~100 sucrose reinforcers/session. Thus, rats learned to reliably self-administer sucrose. **(A)** Average (± SEM) active lever presses, inactive lever presses, and reinforcers earned during fixed ratio 1 (FR1) and fixed ratio 3 (FR3) sessions. Rats in Experiment 1 received 5 sessions, whereas those in Experiment 2 received 4 sessions. Data are collapsed across Experiments 1 and 2. Average (± SEM) **(B)** active and inactive lever presses, **(C)** reinforcers earned, and **(D)** breakpoint achieved under a progressive ratio schedule of sucrose reinforcement (i.e., last total number of lever presses for sucrose before session end) in Experiment 2. *Active lever presses > inactive lever presses across sessions. *n* = 22 females, 23 males.

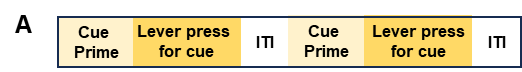

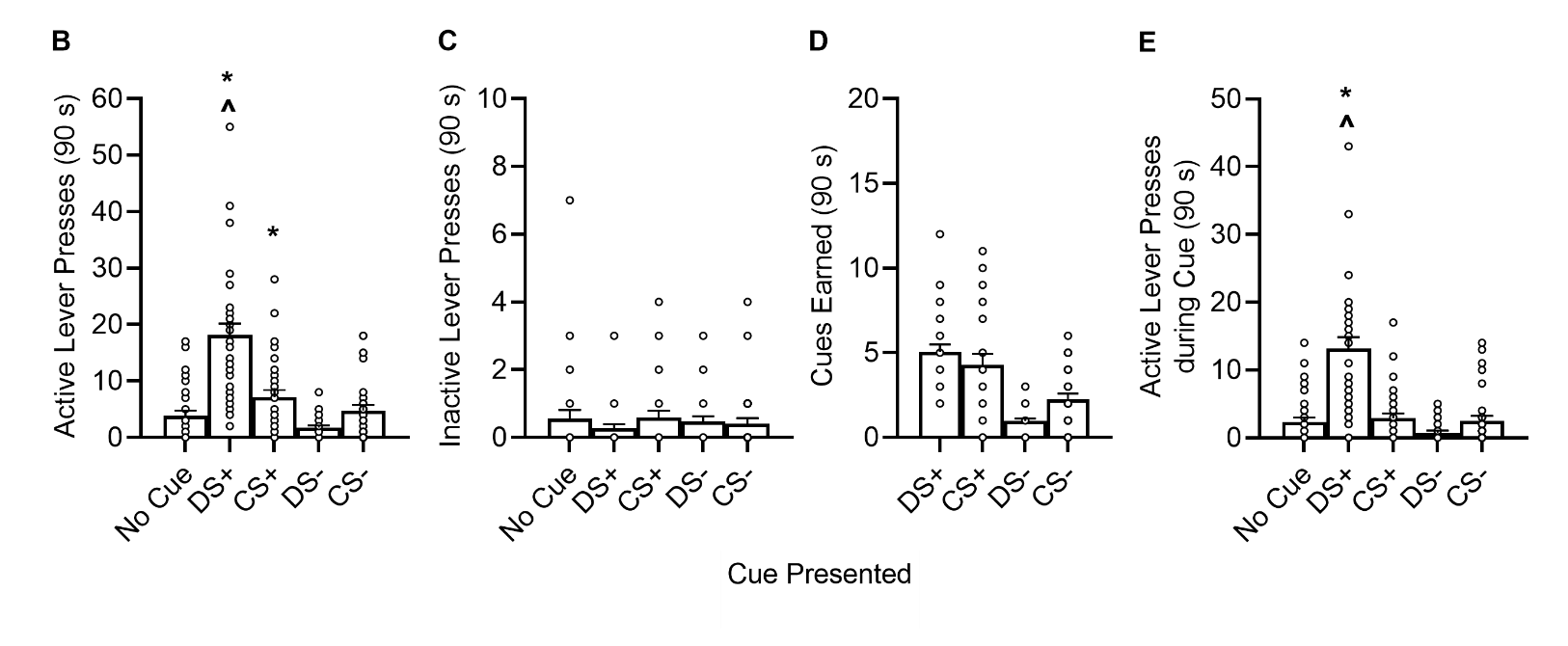

**Supplementary Figure 3. Instrumental responding to obtain presentations of sucrose-associated cues in Experiment 1. (A)** Schematic of the conditioned reinforcement test during which rats could lever press to obtain cue presentations under a Fixed Ratio 1 schedule of reinforcement. All cue conditions were tested in a single session, and each cue condition was tested in 3 x 30-s trials. Average (± SEM) **(B)** active lever presses made to obtain each cue, **(C)** inactive lever presses, **(D)** cue presentations earned, and **(E)** active lever presses made during earned cue presentations. * vs. No Cue. ^ vs. CS+. DS+, discriminative stimulus previously signaling sucrose availability. DS-, discriminative stimulus previously signaling sucrose unavailability. CS+, conditioned stimulus paired with sucrose delivery. CS-, conditioned stimulus never paired with sucrose delivery. ITI, intertrial interval. *n* = 32

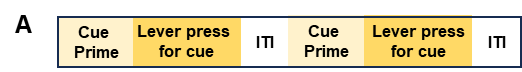

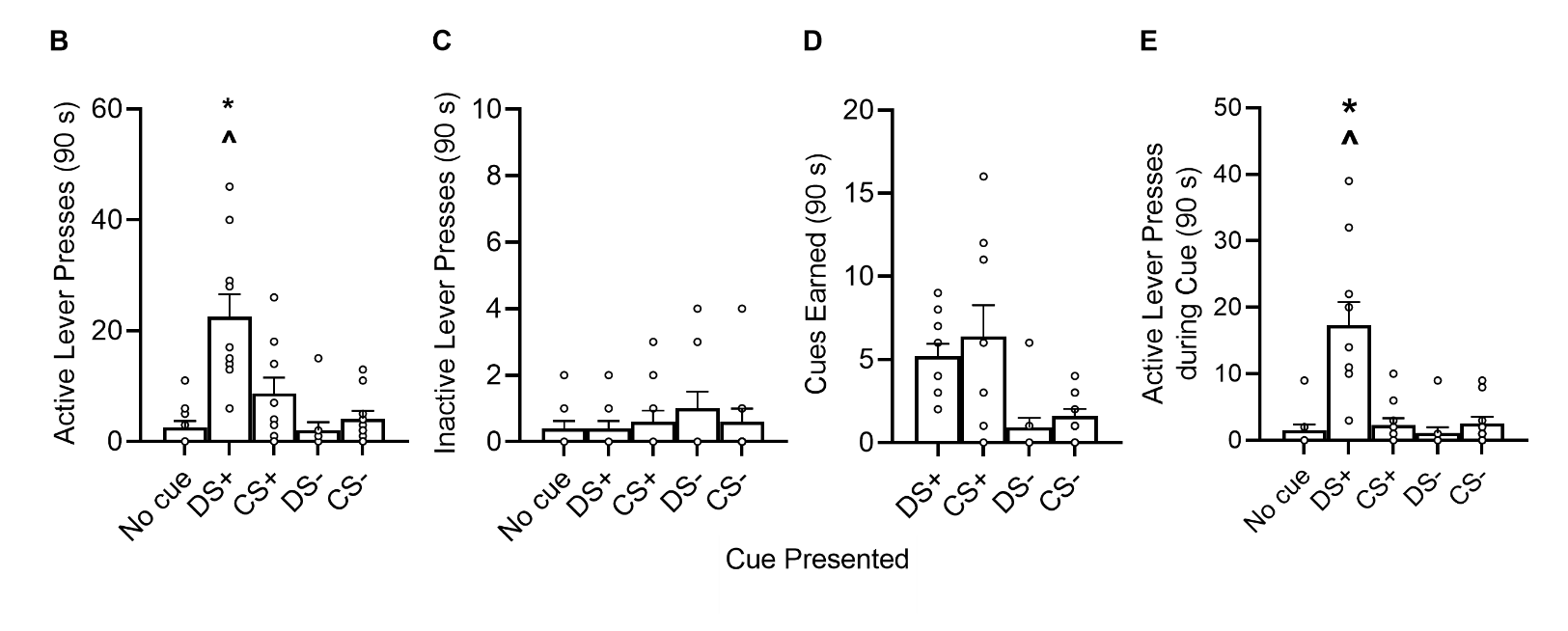

**Supplementary Figure 4. Instrumental responding to obtain presentations sucrose-associated cues in Experiment 2. (A)** Schematic of the conditioned reinforcement test during which pressing on the active lever produced cue presentation under a Fixed Ratio 1 schedule of reinforcement. All cue conditions were tested in a single session, and each cue condition was tested in 3 x 30-s trials. Average (± SEM) **(B)** active lever presses made to obtain the cue, **(C)** inactive lever presses, **(D)** cue presentations earned, and **(E)** active lever presses made during earned cue presentations. * vs. No Cue. ^ vs. CS+. DS+, discriminative stimulus previously signaling sucrose availability. DS-, discriminative stimulus previously signaling sucrose unavailability. CS+, conditioned stimulus paired with sucrose delivery. CS-, conditioned stimulus never paired with sucrose delivery. ITI, intertrial interval. *n* = 10.

**Supplementary Table 1.** Comparing responding in rats with versus without photostimulation-induced seizures in Experiment 1.

|  | **Test statistic** | ***p* value** |
| --- | --- | --- |
| **Discrimination retraining post seizure** | | |
| **ChR2-seizure vs. ChR2-non-seizure** |  |  |
| Session 1: DS+ Ratio | *t*_(17)_ = 0.11 | 0.92 |
| Session 1: DS- Ratio | *t*_(17)_ = 0.11 | 0.92 |
| Session 2: DS+ Ratio | *t*_(17)_ = 1.49 | 0.15 |
| Session 2: DS- Ratio | *t*_(17)_ = 1.49 | 0.15 |
| Session 1: Reinforcers earned | *t*_(13.45)_ = 0.24 | 0.81 |
| Session 2: Reinforcers earned | *t*_(12.26)_ = 0.18 | 0.86 |
| **ChR2-seizure vs. eYFP** |  |  |
| Session 1: DS+ Ratio | *t*_(20)_ = 1.96 | 0.07 |
| Session 1: DS- Ratio | *t*_(20)_ =1.96 | 0.07 |
| Session 2: DS+ Ratio | *t*_(20)_ =1.73 | 0.10 |
| Session 2: DS- Ratio | *t*_(20)_ =1.75 | 0.10 |
| Session 1: Reinforcers earned | *t*_(14.08)_ = 1.21 | 0.25 |
| Session 2: Reinforcers earned | *t*_(20)_ =1.38 | 0.18 |
| **Cue-evoked seeking tests** | | |
| **ChR2-seizure vs. ChR2-non-seizure** |  |  |
| DS+ active lever presses | *t*_(9.44)_ = 1.91 | 0.09 |
| CS+ active lever presses | *t*_(10)_ = 1.16 | 0.28 |
| DS- active lever presses | *z* = -0.85 | 0.76 |
| No cue active lever presses | *z* = -0.27 | 0.88 |
| CS- active lever presses | *t*_(4.53)_ = 1.10 | 0.33 |
| **ChR2-seizure vs. eYFP** |  |  |
| DS+ active lever presses | *z* = -1.33 | 0.19 |
| CS+ active lever presses | *t*_(23)_ = 0.80 | 0.44 |
| DS- active lever presses | *z* = -2.21 | 0.21 |
| No cue active lever presses | *z* = -0.29 | 0.82 |
| CS- active lever presses | *z* = -0.42 | 0.72 |
| **Conditioned reinforcement tests** | | |
| **ChR2-seizure vs. ChR2-non-seizure** |  |  |
| DS+ active lever presses | *t*_(10)_ = -0.12 | 0.90 |
| CS+ active lever presses | *t*_(10)_ = 0.37 | 0.72 |
| DS- active lever presses | *z* = -1.22 | 0.27 |
| No cue active lever presses | *z* = -0.92 | 0.43 |
| CS- active lever presses | *t*_(10)_ = 1.89 | 0.09 |
| **ChR2-seizure vs. eYFP** |  |  |
| DS+ active lever presses | *z* = -0.10 | 0.92 |
| CS+ active lever presses | *z* = - 0.82 | 0.45 |
| DS- active lever presses | *z* = -1.81 | 0.08 |
| No cue active lever presses | *z* = 0.35 | 0.77 |
| CS- active lever presses | *z* = -0.69 | 0.53 |

**Supplementary Table 2.** Comparing responding in females and males across phases in Experiment 1.

|  | **Test statistic** | ***p* value** |
| --- | --- | --- |
| **Sucrose self-administration** | | |
| **Session 1** |  |  |
| Active lever presses | *z* = -0.07 | 0.97 |
| Reinforcers earned | *z* = -1.32 | 0.54 |
| **Session 4** |  |  |
| Active lever presses | *z* = -0.61 | 0.72 |
| Reinforcers earned | *z* = -0.93 | 0.77 |
| **Discrimination training** | | |
| **Session 1** |  |  |
| DS+ ratios | *t*_(20)_ = 1.29 | 0.21 |
| Reinforcers earned | *t*_(20)_ = 0.99 | 0.33 |
| **Session 20** |  |  |
| DS+ ratios | *t*_(20)_ = 0.46 | 0.65 |
| Reinforcers earned | *t*_(20)_ = 2.28 | 0.15 |
| **Cue-evoked seeking test: Active lever presses** | | |
| **Control group** |  |  |
| DS+ | *z* = -1.70 | 0.09 |
| CS+ | *z* = -0.40 | 0.71 |
| DS- | *z* = -0.38 | 0.74 |
| CS- | *z* = -0.00 | 1.00 |
| No Cue | *z* = -0.69 | 0.53 |
| **Stimulation group** |  |  |
| DS+ | *z* = -0.61 | 0.55 |
| CS+ | *z* = -0.93 | 0.37 |
| DS- | *z* = -0.29 | 0.79 |
| CS- | *z* = -0.34 | 0.74 |
| No Cue | *z* = -0.19 | 0.85 |
| **Conditioned reinforcement test: Active lever presses** | | |
| **Control group** |  |  |
| DS+ | *t*_(10)_ = 0.40 | 0.19 |
| CS+ | *t*_(10)_ = 0.19 | 0.85 |
| **Stimulation group** |  |  |
| DS+ | *z* = -0.90 | 0.43 |
| CS+ | *z* = -0.90 | 0.43 |

**Supplementary Table 3.** Comparing responding in females and males across phases in Experiment 2.

|  | **Test statistic** | ***p* value** |
| --- | --- | --- |
| **Sucrose self-administration** | | |
| **Session 1** |  |  |
| Active lever presses | *t*_(21)_ = -2.23 | 0.07 |
| Reinforcers earned | *z* = -2.67 | 0.08 |
| **Session 5** |  |  |
| Active lever presses | *z* = -0.45 | 0.69 |
| Reinforcers earned | *z* = -1.28 | 0.34 |
| **Discrimination training** | | |
| **Session 1** |  |  |
| DS+ ratios | *t*_(21)_ = -0.36 | 0.72 |
| Reinforcers earned | *t*_(21)_ = -0.36 | 0.72 |
| **Session 22** |  |  |
| DS+ ratios | *t*_(21)_ = -3.49 | 0.09 |
| Reinforcers earned | *t*_(21)_ = -3.75 | 0.03* |
| **Cue-evoked seeking test: Active lever presses** | | |
| **Control group** |  |  |
| DS+ | *t*_(31)_ = 0.95 | 0.35 |
| CS+ | *z* = 0.94 | 0.36 |
| DS- | *z* = -1.13 | 0.31 |
| CS- | *z* = -2.21 | 0.14 |
| No Cue | *z* = -0.44 | 0.71 |
| **Stimulation group** |  |  |
| DS+ | *z* = -1.28 | 0.34 |
| CS+ | *z* = 0.00 | 1.00 |
| DS- | *z* = -1.28 | 0.28 |
| CS- | *z* = -0.22 | 0.83 |
| No Cue | *z* = -0.24 | 0.83 |
| **Conditioned reinforcement test: Active lever presses** | | |
| **Control group** |  |  |
| DS+ | *t*_(8)_ = 0.75 | 0.48 |
| CS+ | *t*_(6)_ = -0.27 | 0.80 |
| **Stimulation group** |  |  |
| DS+ | *z* = -0.30 | 0.83 |
| CS+ | *t*_(6)_ = -0.10 | 0.93 |

**Supplementary Table 4.** Correlations between Pavlovian conditioned approach phenotype and responding during cue-evoked sucrose seeking and conditioned reinforcement tests in Experiment 1.

| **Cue-evoked sucrose seeking test** | | | | | | | | | | | | | | | | | | | | | | | | | | |
| --- | --- | --- | --- | --- | --- | --- | --- | --- | --- | --- | --- | --- | --- | --- | --- | --- | --- | --- | --- | --- | --- | --- | --- | --- | --- | --- |
|  | | **Active lever presses** | | | | | | | | | | | |  | | **Magazine entries** | | | | | | | | | | |
|  | | DS+ | | CS+ | | | DS- | | CS- | | | No cue | |  | | DS+ | | | CS+ | | DS- | | CS- | | No cue | |
| ***r*** | | -0.21 | | -0.16 | | | -0.12 | | 0.04 | | | 0.35 | |  | | 0.05 | | | -0.06 | | -0.08 | | 0.26 | | 0.22 | |
| ***p*** | | 0.25 | | 0.37 | | | 0.51 | | 0.81 | | | 0.24 | |  | | 0.80 | | | 0.74 | | 0.66 | | 0.25 | | 0.23 | |
| **Conditioned reinforcement test** | | | | | | | | | | | | | | | | | | | | | | | | | | |
|  | **Active lever presses** | | | | | | | |  | **Cues earned** | | | | | | | |  | | **Magazine entries** | | | | | | |
|  | DS+ | | CS+ | | DS- | CS- | | No  cue |  | DS+ | CS+ | | DS- | | CS- | | No  cue |  | | DS+ | | CS+ | DS- | CS- | | No  cue |
| ***r*** | -0.38 | | -0.04 | | -0.22 | -0.11 | | -0.22 |  | -0.22 | -0.05 | | -0.12 | | -0.05 | | -0.11 |  | | -0.24 | | -0.06 | -0.10 | 0.27 | | -0.22 |
| ***p*** | 0.13 | | 0.83 | | 0.23 | 0.54 | | 0.24 |  | 0.22 | 0.78 | | 0.53 | | 0.78 | | 0.56 |  | | 0.19 | | 0.74 | 0.59 | 0.14 | | 0.23 |

**Supplementary Table 5.** Correlations between Pavlovian conditioned approach phenotype and responding during cue-evoked sucrose seeking and conditioned reinforcement tests in Experiment 2.

| **Cue-evoked seeking test** | | | | | | | | | | | | | | | | | | | | | | | | | | |
| --- | --- | --- | --- | --- | --- | --- | --- | --- | --- | --- | --- | --- | --- | --- | --- | --- | --- | --- | --- | --- | --- | --- | --- | --- | --- | --- |
|  | | **Active lever presses** | | | | | | | | | | | |  | | **Magazine entries** | | | | | | | | | | |
|  | | DS+ | | CS+ | | | DS- | | CS- | | | No cue | |  | | DS+ | | | CS+ | | DS- | | CS- | | No cue | |
| ***r*** | | 0.01 | | -0.01 | | | 0.35 | | 0.01 | | | -0.15 | |  | | -0.13 | | | -0.21 | | 0.12 | | 0.13 | | -0.23 | |
| ***p*** | | 0.99 | | 0.96 | | | 0.23 | | 0.94 | | | 0.41 | |  | | 0.50 | | | 0.24 | | 0.50 | | 0.47 | | 0.20 | |
| **Conditioned reinforcement test** | | | | | | | | | | | | | | | | | | | | | | | | | | |
|  | **Active lever presses** | | | | | | | |  | **Cues earned** | | | | | | | |  | | **Magazine entries** | | | | | | |
|  | DS+ | | CS+ | | DS- | CS- | | No  cue |  | DS+ | CS+ | | DS- | | CS- | | No  cue |  | | DS+ | | CS+ | DS- | CS- | | No  cue |
| ***r*** | 0.17 | | -0.12 | | 0.34 | 0.13 | | 0.21 |  | 0.12 | -0.05 | | 0.24 | | 0.14 | | 0.11 |  | | -0.25 | | 0.28 | -0.18 | -0.59 | | -0.29 |
| ***p*** | 0.64 | | 0.77 | | 0.34 | 0.72 | | 0.57 |  | 0.73 | 0.88 | | 0.51 | | 0.71 | | 0.76 |  | | 0.48 | | 0.44 | 0.63 | 0.07 | | 0.42 |

**Supplementary Table 6.** Correlations between breakpoint for sucrose under a progressive ratio schedule of reinforcement and behavioural responding in Experiment 2.

| **Cue-evoked seeking test** | | | | | | | | | | | | | | | | | | | | |
| --- | --- | --- | --- | --- | --- | --- | --- | --- | --- | --- | --- | --- | --- | --- | --- | --- | --- | --- | --- | --- |
|  | | **Active lever presses** | | | | | | | | |  | | **Magazine entries** | | | | | | | |
|  | | DS+ | CS+ | | DS- | | CS- | | No cue | |  | | DS+ | CS+ | | DS- | | CS- | | No cue |
| ***r*** | | -0.06 | 0.08 | | 0.06 | | 0.08 | | 0.02 | |  | | 0.30 | -0.11 | | -0.26 | | 0.32 | | 0.09 |
| ***p*** | | 0.74 | 0.64 | | 0.75 | | 0.65 | | 0.91 | |  | | 0.05 | 0.50 | | 0.09 | | 0.14 | | 0.56 |
| **Conditioned reinforcement test** | | | | | | | | | | | | | | | | | | | | |
|  | **DS+** | | | | | | | | |  | | **CS+** | | | | | | | | |
|  | Active lever presses | | | DS+ earned | | Active lever presses during DS+ | | Magazine entries | |  | | Active lever presses | | | CS+ earned | | Active lever presses during CS+ | | Magazine entries | |
| ***r*** | -0.32 | | | 0.04 | | -0.22 | | 0.37 | |  | | 0.45 | | | -0.03 | | 0.46 | | 0.23 | |
| ***p*** | 0.37 | | | 0.91 | | 0.47 | | 0.26 | |  | | 0.26 | | | 0.94 | | 0.15 | | 0.55 | |
